# Causal relationship between the right auditory cortex and speech-evoked envelope-following response: Evidence from combined transcranial stimulation and electroencephalography

**DOI:** 10.1101/2020.03.10.985564

**Authors:** Guangting Mai, Peter Howell

## Abstract

Speech-evoked envelope-following response (EFR) reflects brain encoding of speech periodicity that serves as a biomarker for pitch and speech perception and various auditory and language disorders. While EFR is thought to originate from the subcortex, recent research illustrated a right-hemispheric cortical contribution to EFR. However, it is unclear whether this contribution is causal. This study aimed to establish this causality by combining transcranial direct current stimulation (tDCS) and measurement of EFR (pre- and post-tDCS) via scalp-recorded electroencephalography (EEG). We applied tDCS over the left and right auditory cortices in right-handed normal-hearing participants and examined whether altering cortical excitability via tDCS causes changes in EFR during monaural listening to speech syllables. We showed significant changes in EFR magnitude when tDCS was applied over the right auditory cortex compared to sham stimulation for the listening ear contralateral to the stimulation site. No such effect was found when tDCS was applied over the left auditory cortex. Crucially, we further observed a hemispheric laterality where after-effect was significantly greater for tDCS applied over the right than the left auditory cortex in the contralateral ear condition. Our finding thus provides the first evidence that validates the causal relationship between the right auditory cortex and EFR.

## Introduction

The speech-evoked frequency-following responses (FFR) are phase-locked neural activities that reflect early processing of periodic features of input speech signals in the human brain (Picton and Aiken, 2008; Coffey et al., 2019). One of the most important FFR components in the central auditory systems is the envelope-following response (EFR) which encodes the periodicity envelopes at fundamental frequency (F_0_) that represents the vocal pitch information (Picton and Aiken, 2008; Coffey et al., 2019). The EFR is associated with various human auditory and language processing. For example, it reflects the neural encoding of linguistic pitch and is stronger in tonal language than non-tonal language speakers (Krishnan et al., 2004, 2005, 2009). It has greater strength in musicians who have better pitch discrimination ability than people without musical training (Musacchia et al., 2007; Wong et al., 2007; Strait et al., 2009; Bidelman et al., 2011). Also, EFR is associated with speech-in-noise perception. Greater EFR magnitudes are associated with better speech recognition ability in noisy environments (Song et al., 2011; Parbery-Clark et al., 2011). EFR also reflects neural plasticity related to fundamental cognitive and physiological processes such as auditory learning (Skoe et al., 2014), changes in arousal (Mai et al., 2019) and attention (Lehmann and Schönwiesner, 2014; Hartmann and Weisz, 2019).

Clinically, EFR is proposed as a potential biomarker for various auditory and language disorders. EFR declines with age (Anderson et al., 2012; Presacco et al., 2016) and can predict word recognition ability during speech-in-noise perception in older adults (Anderson et al., 2011; Fujihira and Shiraishi, 2015; Mai et al., 2018). This indicates that degradations to EFR could potentially explain the increased speech-in-noise difficulty experienced during aging. EFRs are also associated with hearing deficits such as cochlear synaptopathy (Encina-Llamas et al., 2019) and auditory processing disorders (Schochat et al., 2017). Furthermore, deficits of EFR often occurred along with functional impairments in learning and cognitive disorders, such as learning difficulties in literacy (Cunningham et al., 2001; Banai et al., 2007; White-Schwoch et al., 2015), dyslexia (Hornickel et al., 2013) and autism (Russo et al., 2008) in children and mild cognitive impairment in older adults (Bidelman et al., 2017).

Because of the relationship between EFR and these fundamental and clinical auditory and language processes, it is essential to understand how different parts of the auditory systems may contribute to EFR. It has long been argued that EFR reflects the encoding of periodicity in the inferior colliculus at the brainstem, which has been proposed as the main neural source of EFR (Chandrasekaran and Kraus, 2010; Bidelman, 2015, 2018). Recent studies, however, have shown cortical contributions to EFR (Coffey et al., 2016, 2017a; Hartmann and Weisz, 2019; Ross et al., 2020). These studies localized the EFR sources along the auditory pathway using magnetoencephalography (MEG) and illustrated significant neural contributions to EFR within the right auditory cortex associated with musical experience, pitch discrimination ability (Coffey et al., 2016), speech-in-noise perception (Coffey et al., 2017a), intermodal attention (Hartmann and Weisz, 2019) and ageing (Ross et al., 2020). Another study found that EFR strength was associated with right, but not left, hemispheric hemodynamic activity in the auditory cortex (Coffey et al., 2017b), consistent with the relative specialization of right auditory cortex for pitch and tonal processing (Zatorre and Berlin, 2001; Patterson et al., 2002; Hyde et al., 2008; Albouy et al., 2013; Cha et al., 2016).

Despite findings that show the potential cortical contribution to EFRs, it is unclear whether the relationship between auditory cortex and EFR is causal and is not simply an observation based on specific source localization techniques or apparent association between cortical activations and EFR. More specifically, it is unclear whether EFR is susceptible to neuroplasticity at the cortical level, e.g., how plastic changes in neural excitability of the auditory cortex may cause changes in EFR. An important next step, therefore, is to determine such causality. One way to achieve this aim is to use neuro-stimulation tools that are known to change cortical excitability and test how EFR may be altered according to these changes. A recent study investigated how EFR obtained via scalp-recorded electroencephalography (EEG) was modulated by inhibiting excitability of the right auditory cortex using repetitive transcranial magnetic stimulation (rTMS) (Lopez-Caballero et al., 2020). This study, however, did not find significant changes in EFR following stimulation compared to sham. The authors argued that the absence of the after-effect may be attributed to ineffectiveness of rTMS to change neural excitability in the auditory sensory region (Lopez-Caballero et al., 2020). Here, we address several critical caveats. First, it is possible that the right auditory cortex contributes to EFR mainly along the contralateral pathway. Stimulation over the right auditory cortex might result in significant changes in EFR when participants listen to sounds from the contralateral ear (i.e., the left ear from which the right auditory cortex receives the majority of its auditory information), rather than the ipsilateral ear (i.e., the right ear). If this is the case, binaural listening during EFR measurements in Lopez-Caballero et al (2020) might smear the rTMS effect dominant in the contralateral ear. Therefore, using paradigms of monaural listening that allow studying ipsilateral/contralateral effects of stimulation should provide better insights into cortical contributions to EFR. Second, although previous research suggested a right-hemispheric cortical contribution to EFR (Coffey et al., 2016, 2017a, 2017b; Hartmann and Weisz, 2019; Ross et al., 2020), a more rigorous design is to include the left auditory cortex as a control stimulation site. This could determine whether cortical contribution is made specifically in the right hemisphere and whether laterality effects occur by comparing after-effects of stimulations over different hemispheres. Finally, EEG has a poor source localization capability to determine the site of occurrence of the after-effects. It is therefore necessary to look into the after-effects at different neural latencies. EFR originates at the auditory brainstem at 5L15 ms after stimulus onset (Chandrasekaran and Kraus, 2010). Recent research has further illustrated that there are cortical EFR activities that peak at 50L60 ms after sound onset (Coffey et al., 2016). Looking into timing characteristics of after-effects could illustrate whether any effects start at the early (brainstem) or later (auditory cortex) auditory centres. Note that ‘cortical contribution’ here we refer to includes not only contributions of cortical EFR, but also corticofugal modulation on the subcortical EFR (Price et al., 2021) even without cortical sources. Examining the timing characteristics could help disentangle whether the cortical contribution is likely made directly at the cortex or through top-down corticofugal modulation on the subcortical level.

Based on the discussions above, the aim of the current study was to establish a causal relationship between the auditory cortex and EFR. Here, we applied transcranial direct current stimulation (tDCS) to alter neural excitability in the left and right auditory cortices. We measured the EFR using scalp-recorded EEG during monaural listening (left and right ears respectively to allow for testing contralaterality) to a repeatedly-presented speech syllable pre- and post-tDCS. We then examined the tDCS after-effects on EFR. tDCS is a non-invasive neuro-stimulation that modulates cortical excitability (Jacobson et al., 2012). By applying direct currents over the scalp, tDCS leads to neural excitation or inhibition in proximal parts of the cortex that last for up to 90 minutes post-stimulation (Nitsche and Paulus, 2001). Previous research has shown that tDCS over auditory cortices can modulate cortical auditory-evoked responses to pitch (recorded via EEG) (Zaehle et al., 2011; Impey and Knott, 2015; Boroda et al., 2020). As for hemispheric laterality, previous studies showed that applying tDCS over the right, compared to the left, auditory cortex can significantly change pitch discrimination performances, supporting the causal role of the right auditory cortex for pitch perception (Mathys et al., 2010; Matsushita et al., 2015, 2021). However, such causality has not been established for neurophysiological signatures like EFR. Should cortical contributions to EFR exist, altering cortical excitability via tDCS should cause consequent changes in EFR. The current study tested the hypothesis that tDCS over the right auditory cortex results in changes in EFR which should occur particularly along the contralateral auditory pathway where participants listen to speech from the left ear. Besides, we further examined the hemispheric laterality by directly comparing stimulations between the two hemispheres and investigated the timing characteristics of the after-effects.

## Materials and Methods

The present study was approved by the UCL Research Ethics Committee and informed consents were obtained from all participants that were recruited for the experiments.

### Participants

Ninety right-handed, normal-hearing participants (18-40 years old; 45 females) were recruited and completed the entire experiment. Two other participants dropped out during the tDCS phase because they felt uncomfortable with the skin sensation when stimulation was applied. All participants are right-handed (Handedness Index (HI) > 40; Oldfield, 1971) and had normal hearing (pure-tone audiometric thresholds (PTA) 25 ≤dB HL within the range of 0.25–6 kHz for both ears). Participants were all non-tonal language (English, Spanish, Portuguese, Polish or Russian) speakers, had no long-term musical training and reported no history of neurological or speech/language disorders. No participants participated in any brain stimulation experiments within the two weeks prior to the present experiment.

Participants were assigned at random to one of five groups, each of which received different types of tDCS. Handedness (HI), age, gender and audiometric thresholds were all matched across the five groups (see *tDCS* in *Experimental design* for details).

### Experimental design

The experimental procedure is summarized in **Figure 1**. EFRs were recorded pre- and post-tDCS during monaural listening to a repeatedly-presented speech syllable to test for any after-effects of tDCS. Details of the EFR recording and tDCS protocols are described below.

**Figure 1.**
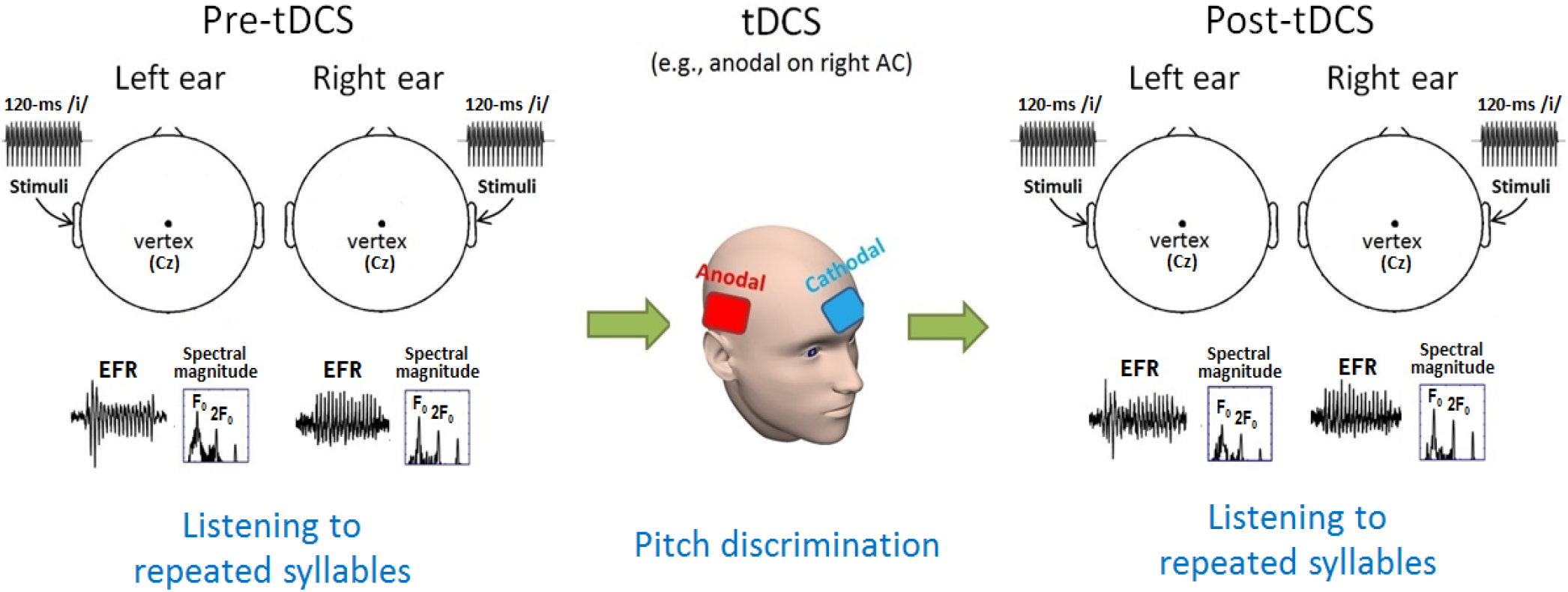
The experimental design. Participants underwent three phases (pre-tDCS, tDCS and post-tDCS). EFRs were measured via scalp EEG at Cz when participants listened to a repeatedly presented syllable /i/ (120 ms long) monaurally pre- and post-tDCS. A pitch discrimination task was performed during the tDCS application over the auditory cortex with the reference electrode placed above the contralateral eyebrow on the forehead.

#### Speech stimulus for the EFR recording

A 120-ms-long syllable /i/ spoken by a male with a static fundamental frequency (F_0_) at 136 Hz was used for the EFR recordings. The reason for using /i/ was that EFR elicited by this stimulus had been proved to be robustly measured in our lab (Mai et al., 2018, 2019). The syllable has a stable amplitude profile across the syllable period except for the 5-ms rising and falling cosine ramps applied at the onset and offset to avoid transients. The waveform of the syllable is shown in **Figure 2A**.

**Figure 2.**
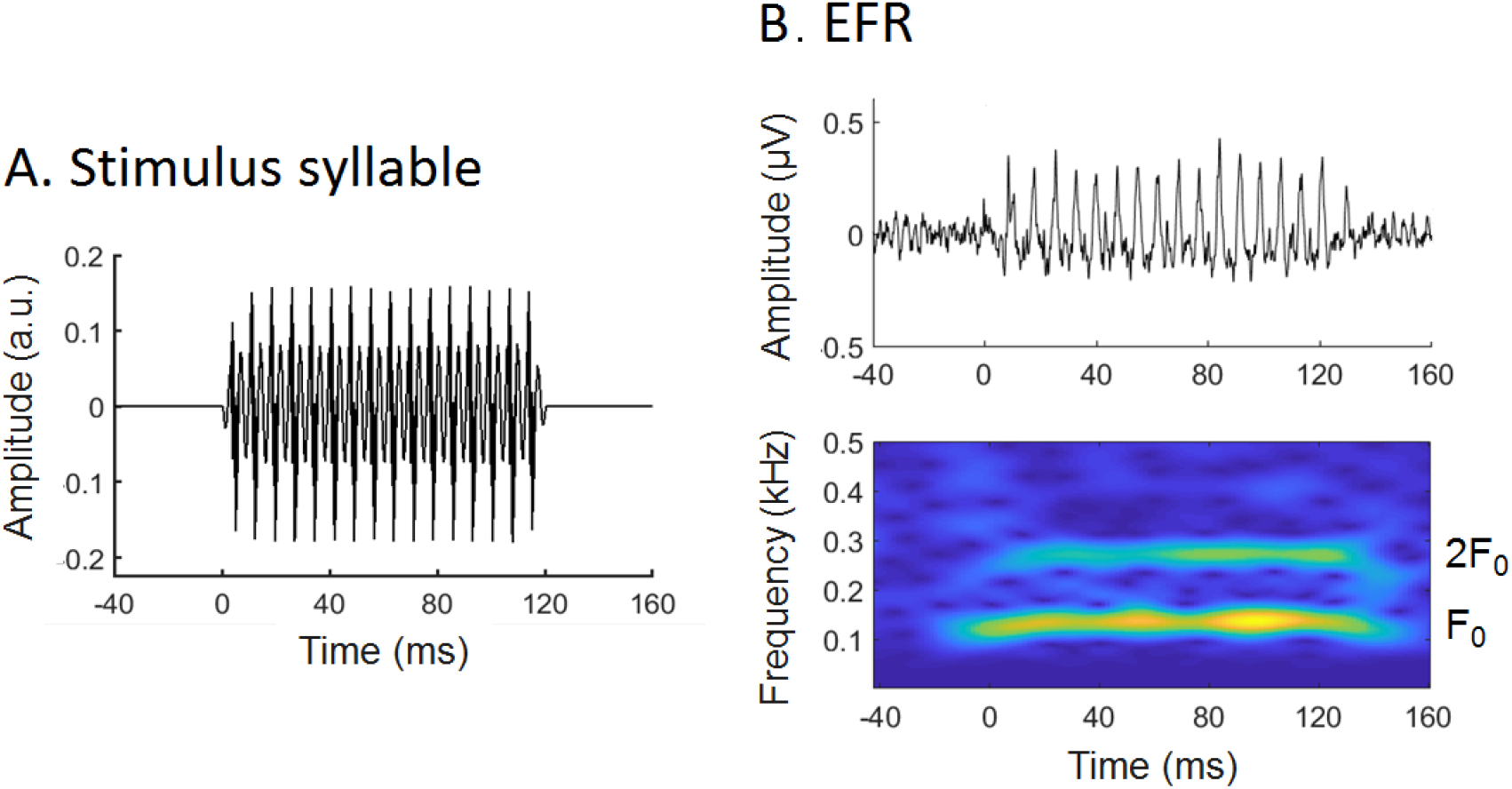
The syllable stimulus for EFR recordings. (A) Temporal waveform of the 120-ms stimulus syllable /i/ with fundamental frequency (F_0_) at 136 Hz. **(B)** A representative sample waveform of EFR (upper) and its spectrogram (lower) showing that the powers are predominated by F_0_ and 2F_0_.

The syllable stimulus was repeatedly presented at ∼4 times per second with inter-stimulus interval (ISI, time interval between offset of one stimulus and offset of the following stimulus) fixed at 120 ms. The stimulus was played monaurally via electromagnetically-shielded inserted earphone (ER-3 insert earphone, Intelligent Hearing Systems, Miami, FL) at 85 dB SPL (excluding ISIs) in each ear (e.g. left ear listening followed by right ear listening or vice versa with the order of ear presentation counterbalanced across participants). For each ear, there were 3,000 sweeps in total with half of the sweeps (i.e., 1,500 sweeps) having positive polarity and the other half having negative polarity. The reason for having two different polarities was to minimize the neural responses to the temporal fine structure information which mainly occur at auditory periphery (Aiken and Picton, 2008) and electrical artefacts by adding the responses to the positive and negative sweeps (see *EEG signal processing*). Sweeps with different polarities were presented in an intermixed order for all participants (set prior to the experiments via pseudo-randomization).

#### EFR recording

EEGs were recorded over participants’ scalps using an ActiveTwo system (Biosemi ActiView, The Netherlands) with sampling rate of 16,384 Hz whilst they listened to the 3,000 sweeps of the syllable stimulus both pre- and post-tDCS. Due to the time constraints for which the EFR recording needed to be completed as soon as possible after finishing the tDCS phase (in order to maintain the offline effect of tDCS), only three active electrodes (Cz, C3 and C4 localized using a standard Biosemi cap) were placed to save time for EEG setups. We only focused on Cz only as it is the standard conventional site used for obtaining robust EFR (Skoe and Kraus, 2010). Bilateral earlobes served as the reference and ground electrodes were CMS and DRL at the parieto-occipital sites. Electrode offsets were always kept within ±35 mV.

Participants were seated comfortably in an armchair in an electromagnetically- and sound-shielded booth. They listened passively to the stimuli whilst keeping their eyes on a fixation cross in the centre of a computer screen. The 3,000 syllable sweeps in each ear were broken into six 2-minute-long blocks (500 sweeps each) with ∼40 second breaks between blocks to minimize fatigue. Participants were required to keep awake and refrain from body and head movements whilst they were listening to the sounds. Participants were instructed to keep awake because our previous study has shown that EFR magnitudes decrease significantly when participants fall sleep (Mai et al., 2019). Allowing participants to sleep could result in varying levels of arousal that affected the EFR in an uncontrolled manner and therefore could cause confounds for quantifying the tDCS after-effects. The EFR recording lasted for ∼30 minutes for both pre- and post-tDCS. The post-tDCS recording was completed within 45 minutes after tDCS was stopped for all participants to ensure that any after-effects of tDCS on EFRs were sustained (Nitsche and Paulus, 2001).

#### PTA

Along with EFR recording, audiometric thresholds (PTA at 0.25, 0.5, 1, 2, 3, 4 and 6 kHz) in both ears were also tested pre- and post-tDCS using a MAICO MA41 Audiometer (MAICO Diagnostics, Germany). These were respectively conducted prior to the pre- and post-tDCS EFR recording to test whether tDCS changed peripheral hearing and whether after-effects on EFR were related to these changes because EFR could be influenced by individual’s audibility (Ananthakrishnan et al., 2016). Seven participants’ post-tDCS PTA data were not recorded (one in the Left-AC Anodal group, two in the Right-AC Anodal group, three in the Right-AC Cathodal group and one in the Sham group; see group allocations in *tDCS*) due to the unavailability of equipment at the time of testing. Other than these absent data, PTAs at all frequencies were within the normal range ( 25 ≤dB HL) both pre- and post-tDCS in both ears.

Subsequent analyses showed baseline (pre-tDCS) PTA and change in PTA (post- vs. pre-tDCS) did not differ significantly between stimulation groups for either ear (see *tDCS* for details). Also no significant association was found between the after-effect on EFR and baseline PTA or change in PTA (see *Results*).

#### tDCS

tDCS was applied over the scalp using a battery-driven direct current stimulator (Magstim HDCStim, UK) with a pair of rubber-surface electrodes (5×5 cm) contained in saline-soaked cotton pads. Centre position of the active electrode was on T7/T8 (according to the 10/20 EEG system) for the left/right auditory cortex. The exact placement of T7/T8 was determined with the help of a standard Biosemi EEG cap. The reference electrode was placed on the forehead above the eyebrow contralateral to the active electrode (see Matsushita et al., 2015; also see **Figure 1**).

Participants were assigned at random to one of the five groups (18 participants per group; single-blinded). The five groups received the following different types of tDCS: (1) anodal stimulation on the left auditory cortex (Left-AC Anodal); (2) cathodal stimulation on the left auditory cortex (Left-AC Cathodal); (3) anodal stimulation on the right auditory cortex (Right- AC Anodal); (4) cathodal stimulation on the right auditory cortex (Right-AC Cathodal); and (5) Sham. There were equal numbers of males and females in each group. Subsequent checks via independent-sample t-tests between groups confirmed that participants’ age, PTA (averaged across 0.25–6 kHz for each ear), changes in PTA (post- vs pre-tDCS for each ear), and handedness (HI) were matched across groups (all *p* > 0.3, False Discovery Rate (FDR) corrected according to multiple (ten) comparisons between the five groups). Matching of these factors is important because age and peripheral hearing could influence EFR strengths (Anderson et al., 2012; Ananthakrishnan et al., 2016), whilst handedness is associated with hemispheric specialization (Carey and Johnston, 2014; Willems et al., 2014). The matching thus minimized possible confounds on any tDCS effects caused by these factors.

tDCS was applied at 1 mA for 25 minutes with the currents ramping up/down for 15 seconds at the stimulation onset/offset. For Sham, actual stimulation was applied for only 30 seconds in total (15 seconds ramping up and down respectively) at the very beginning of the 25-minute period. This created the usual sensations associated with tDCS but without actual stimulation during the remainder of the run. The electrode configuration for each participant in the Sham group was randomly chosen from (1)–(4) except that it was ensured that the active electrode was positioned either on the left or on the right auditory cortex for an equal number of participants. Compared to other stimulation techniques like TMS which emits loud clicking sound during real stimulation, tDCS is silent, hence avoiding acoustic confounds on blinding participants. After the experiment, most participants, including those in the Sham group, orally reported that they believed they had received real stimulation. All experimental sessions were conducted during daytime (mornings or early afternoons) and all participants reported that they had at least 6 hrs sleep the night before which ensured adequate cortical plasticity triggered by tDCS (Salehinejad et al., 2019).

During tDCS, participants completed a pitch discrimination task while they listened to sound stimuli over a loudspeaker 1 metre in front of them in the same sound-shielded booth used for the EFR recordings. Three short complex tones (400 ms) calibrated at 75 dB SL at the 1 metre position were presented on each trial. The task was an ‘ABX’ task. In each trial, two tones ‘A’ and ‘B’ with different F_0_ (one of which had the same F_0_ at 136 Hz as for the syllable used in the EFR recording) were played consecutively followed by a third tone ‘X’ randomly selected from ‘A’ or ‘B’. Participants had to identify whether ‘X’ was the same as ‘A’ or ‘B’. They gave their best guess if unsure of the answer. The process followed a ‘2-down, 1-up’ adaptive procedure: F_0_ difference between ‘A’ and ‘B’ decreased by times following two √2 consecutive correct trials and increased by times following an incorrect trial. No feedback √2v about response accuracy was provided. Half-minute breaks were taken every 4 minutes. The total number of completed trials was between 160 and 200 based on each participant’s own pace. This task was included during tDCS because tDCS preferentially modulates neural networks that are currently active (Reato et al., 2010; Ranieri et al., 2012; Bikson and Rahman, 2013). Concurrent tDCS and the pitch discrimination task could therefore maintain auditory cortical activity during neuro-stimulation, hence maximizing the effect of tDCS on neural excitability.

### EEG Signal processing

All EEG signal processing was conducted via Matlab R2017a (The Mathworks).

#### Pre-processing

As mentioned, the EFR was captured from Cz. The EEG signals were first re-referenced to the bilateral earlobes and bandpass filtered between 90 and 4000 Hz using a 2nd-order zero-phase Butterworth filter. The filtered signals were then segmented and baseline-corrected (subtracting the average of the 50-ms pre-stimulus period) for each sweep. Sweeps that exceeded ±25 µV were rejected to minimize movement artefacts. The resultant rejection rates were less than 2.5% averaged across participants in all cases (pre- and post-tDCS in the five stimulation groups for both left and right ear conditions).

#### EFR magnitudes

Frequency-following responses (FFRs) with the positive and negative polarities (FFR_Pos_ and FFR_Neg_) were first obtained by temporally averaging the pre-processed signals across sweeps with the respective polarities. EFR was measured as the response to the envelopes of F_0_ and its harmonics by adding FFR_Pos_ and FFR_Neg_ (Aiken and Picton, 2008). The addition of responses to the syllables with two polarities minimized the responses to temporal fine structures and cochlear microphonics, so that purer responses to envelopes were obtained to reflect the encoding of speech periodicity (Aiken and Picton, 2008). A representative sample of EFR (waveform and spectrogram) is shown as **Figure 2B**. Here, EFR_F0_ and EFR_2F0_ (EFR at F_0_ and its 2nd harmonic, 2F_0_) that dominate the power of EFR were focused on. In contrast to higher harmonics (≥3) of EFR that may reflect distortion products resulting from non-linear auditory response on the basilar membrane, EFR_F0_ and EFR_2F0_ reflect neural phase-locking to sound periodicity in the central auditory systems (Smalt et al., 2012). Whilst it is expected that EFR_F0_ plays the major role in the phase-locking, EFR_2F0_ also makes contributions (Aiken and Picton, 2008) because of the non-sinusoidal characteristics of speech periodicity (Holmberg et al., 1988; also see discussions in Smalt et al., 2012). Magnitudes of EFR_F0_ and EFR_2F0_ were then measured following the procedure as follows:

(1) Optimal neural lags relative to the syllable onset that generated the maximal stimulus-to-response correlations (Krizman and Kraus, 2019) were first obtained for EFR. Specifically, both the syllable stimulus and EFR waveform were bandpass filtered at 126–146 and 262–282 Hz (i.e., 20 Hz bandwidth with centre frequencies at F_0_ and 2F_0_) using a 2nd-order zero-phase Butterworth filter. 100 ms pre- and post-stimulus periods were included during filtering to prevent filtering-induced boundary artefacts from contaminating the waveform at the stimulus period. Cross-correlations between the filtered stimulus and EFR were then conducted over a range of time delays (EFR lagged behind the stimulus) for the stimulus period at F_0_ and 2F_0_, respectively. This range was set at 6–16 ms, for which 5–15 ms are latencies when EFR starts to occur in the brainstem (Chandrasekaran and Kraus, 2010) with an additional 1 ms for sound transmission through the earphone plastic tube to the cochlea. The time delay that corresponded to the maximum absolute correlation value was treated as the optimal neural lag.
(2) Magnitudes of EFR were then measured (for EFR_F0_ and EFR_2F0_ separately). The EFR waveform was windowed with a 5-ms rising and falling cosine ramp at the onset and offset. Note that the waveform was the one after pre-processing but prior to step (1), i.e., the bandpass filtering described in step (1) was for determining the optimal neural lags only, but not for measuring EFR magnitudes here in step (2). The window was set to lag behind the stimulus with the optimal neural lag obtained in the previous step (for EFR_F0_ and EFR_2F0_, respectively). The windowed EFR waveform was then zero-padded to 1 second to allow for 1 Hz frequency resolution and the log-transformed FFT-power spectrum (10*log_10_[power]) was measured. EFR_F0_ and EFR_2F0_ magnitudes were taken as the powers centered at F_0_ and 2F_0_ (averaged across 136 ± 2 Hz and 272 ± 2 Hz), respectively.

### Statistical analyses

All statistical analyses were conducted using SPSS Statistics 26.0 (IBM).

#### Baseline characteristics

Before testing the after-effects of tDCS, statistics were first conducted to check whether baseline (pre-tDCS) EFR characteristics were matched across stimulation groups. Linear mixed-effect regressions were conducted based on the restricted maximum likelihood (REML) approach. Baseline EFR magnitude and optimal neural lag were used as dependent variables. Stimulation Group (the five stimulation groups), Ear (left vs right ear) and Harmonic (F_0_ vs 2F_0_) were fixed-effect factors, and Participant was the random-effect factor. The covariance matrix type was chosen among commonly-used structures (first-order auto-regression, compound symmetry, diagonal, scaled identity, Toeplitz and unstructured) to generate the lowest Bayesian Information Criterion (BIC) values (i.e., best goodness of fit). The degrees of freedom were estimated via Satterthwaite approximation. Post-hoc analyses were planned following significant interactions or main effects (Maxwell and Delaney, 2004). Specifically, when there was a significant interaction, the whole dataset was split by the interactive factors and additional linear mixed-effect regressions were conducted at the respective levels within these factors. This procedure was repeated until a significant main effect was found. Pair-wise comparisons were finally conducted between the underlying levels following a main effect (unless there were only two levels within the factor for the main effect, since in such cases the main effect already informed the significant difference between levels). *P* values for pairwise comparisons were corrected via False Discovery Rate (FDR) according to the multiple number of comparisons.

#### After-effects across stimulation groups and ears

Analyses were then conducted to investigate how after-effects of tDCS differed across stimulation groups and ears. Although we mainly focused on the after-effects for EFR magnitude, we also tested after-effects for the neural lag, because previous research also found that there is a potential association between the EFR neural lag and auditory cortical activities (Coffey et al., 2017b).

Linear mixed-effect regressions using the same fixed- (Stimulation Group, Ear and Harmonic) and random-effect (Participant) factors as for the baseline EFR were conducted. The dependent variables were the after-effects of tDCS (i.e., differences in EFR magnitude and neural lag between post- and pre-tDCS). Here, we normalized the after-effects to z-values according to Sham (Grami et al., 2021) before the mixed-effect regressions were conducted. The z-values were obtained as follows:

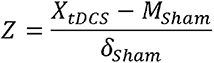

where *X_tDCS_* denotes the raw after-effects (either real tDCS or Sham), *M_Sham_* and δ*_Sham_* denote the mean and standard deviation of the raw after-effects across participants in the Sham group. Note that z-values in the left and right ear conditions were calculated separately, i.e., z-values in the left ear condition were calculated according to Sham in the left ear condition whilst z-values in the right ear condition were calculated according to Sham in the right ear condition. Compared to the raw values of the after-effects, the z-values accounts for the Sham effect which normalizes the data to better reflect after-effects of the real tDCS (Grami et al., 2021).

Additional analyses were conducted using several potential confounding factors respectively as a fixed-effect covariate in the linear mixed-effect regressions. These factors include age, PTA (pre-tDCS), change in PTA (post- vs pre-tDCS) and handedness (HI) (all mean-centred; missing data for the change in PTA were set as zero). Although all these factors were matched across stimulation groups (see *Experimental design*), such additional analyses were conducted to confirm that they did not contribute statistically to the after-effects.

#### Laterality and contralaterality of after-effects

The linear mixed-effect regressions described in the previous section tested the after-effects across stimulation groups and ears. Importantly, they informed how after-effects of real tDCS over the auditory cortex differed significantly from Sham. However, they were unable to directly test the main effect of the stimulated hemisphere (i.e., left vs right auditory cortex) which reflects hemispheric laterality of cortical contributions and how laterality may depend on whether the listening ear was ipsilateral/contralateral to the stimulated site. This is because data of the Sham group was included in the analyses and there was no definitive correspondence of Sham to specific stimulated hemispheres in the current experimental design (see *Experimental design*).

Therefore, a further linear mixed-effect regression was conducted by excluding data in the Sham group to test the laterality and contralaterality of the after-effects. Z-value of the after-effect was used as the dependent variable that accounted for the Sham effect. We first conducted a preliminary analysis using the following four fixed-effect factors: Stimulated Hemisphere (the left vs. right auditory cortex), Contralaterality (the ipsilateral vs. contralateral ear to the stimulated hemisphere), Stimulation Type (anodal vs. cathodal) and Harmonic (F_0_ vs. 2F_0_). Participant was the random-effect factor. We found that neither Stimulation Type nor Harmonic show significant main effects or interactions with other factors (all *p* > 0.1), indicating that effects of these two factors were negligible. Therefore, in order to reduce the analysis complexity and highlight the key effects, data of participants who received anodal and cathodal tDCS were grouped together and the two harmonics were collapsed. The linear mixed-effect regression was then conducted using only Stimulated Hemisphere and Contralaterality as fixed-effect factors and Participant as the random-effect factor. Post-hoc analyses were conducted following significant interactions/main effects.

Note that this linear mixed-effect regression was conducted only for EFR magnitude, but not for the neural lag. This is because significant after-effects of real tDCS compared to Sham were only found for EFR magnitude (according to analyses in the previous section, see *Results*), following which laterality and contralaterality were further tested.

#### Timing characteristics of significant after-effects

Analyses were conducted to study when significant after-effects on EFR magnitudes started to emerge by looking into variations of magnitudes across multiple time frames. Instead of measuring FFT power used for the linear mixed-effect regressions, we measured the power of temporal amplitude profile of the EFR waveform. This is because, empirically, the length of each time frame should be at least two F_0_ cycles (14.7 ms) in order to accurately measure FFT power at F_0_ (Liu and Morgan, 2006). In contrast, a shorter length for each frame is needed for measuring power of amplitude profile which results in better temporal resolution. This is important because the emergence of subcortical/brainstem responses occur within a very narrow time window (5–15 ms after stimulus onset), hence relatively fine-grained temporal resolution is needed to capture detailed magnitude variations so that emergence at subcortical and cortical responses can be disentangled.

Waveforms of EFR were first band-pass filtered using a 2nd-order zero-phase Butterworth filter at 100–300 Hz (for F_0_ plus 2F_0_) and 100–200 Hz (for F_0_). The filtered waveforms were then Hilbert-transformed to obtain the Hilbert envelope as the temporal amplitude profile. The profile was then segmented into successive time frames time-locked to the stimulus onset. The length of each frame was set at 7.4 ms (corresponding to one cycle of F_0_). We focused on the first seven frames up till ∼50 ms (Frame 7 ranged at 44.4–51.8 ms after stimulus onset) that cover the latencies from brainstem to the primary auditory cortex (see Coffey et al., 2016) at which after-effects could emerge. Magnitudes were measured as the log-power of the amplitude profile within each frame and were z-normalized according to Sham as conducted for the linear-mixed effect regressions.

The following two comparisons were then conducted via independent-sample t-tests for the magnitudes in each frame: (a) tDCS over the right auditory cortex (Right-AC Anodal and Cathodal) vs. Sham in the left ear condition (Comparison (a)); and (b) tDCS over the right (Right-AC Anodal and Cathodal) vs. left auditory cortex (Left-AC Anodal and Cathodal) in their respective contralateral ear conditions (Comparison (b)). These comparisons were chosen because they were found to be significant according to results of the linear mixed-effect regressions (see *Results*). For simplicity, data for anodal and cathodal stimulation were grouped together for the t-tests, because no significant interactions involved with Stimulation Type (anodal vs. cathodal) were found in the previous linear mixed-effect regressions and significant after-effects of Right-AC Anodal and Cathodal led to the same direction of changes (see *Results*). The comparisons were conducted via combining the two harmonics (F_0_ plus 2F_0_) due to the lack of interactions involved with Harmonic in the previous linear mixed-effect regressions. Comparisons were also conducted for the F_0_ only because the upper frequency limit of cortical EFR is within the range of vocal pitch (< 200 Hz, Guo et al., 2021). It is thus meaningful to look specifically into whether after-effects at F_0_ alone would start at a relatively later time frame at the cortical level. *P* values of the t-tests were all FDR-corrected according to the total number of steps (i.e., seven).

#### Pitch discrimination performances

Pitch discrimination was performed during the tDCS application (see *Experimental design*) and the change in discrimination performances was calculated for each participant. On each trial, pitch difference of the two complex tones (‘A’ and ‘B’) was recorded. The initial and final discrimination thresholds were measured as the geometric means of the pitch differences across trials with the first ten and final ten reversals, respectively. The change in discrimination performances was then measured as the final threshold divided by the initial threshold.

We tested how changes in pitch discrimination differed across stimulation groups. A linear mixed-effect regression was conducted using the change in pitch discrimination performances as the dependent variable, Stimulation Group as the fixed-effect factor and Participant as the random-effect factor. We also tested how changes in pitch discrimination were associated with changes in EFRs. We conducted Pearson’s correlations between the changes in performances and after-effects on EFR magnitudes in the respective stimulation groups. *P* values of correlations were FDR-corrected according to the total number of correlations conducted.

## Results

### Baseline characteristics

Linear mixed-effect regressions were conducted for the baseline (pre-tDCS) EFR magnitude and neural lag.

For the EFR magnitude, there were significant main effects of Ear (F(1, 116.140) = 12.978, *p* < 0.001, greater magnitude in the left than the right ear with a small to medium effect size (Cohen’s *d* = 0.376)) (**Figure 3**) and Harmonic (F(1, 90.631) = 67.029, *p* < 10^-11^, greater magnitude at F_0_ than at 2F_0_ with a large effect size (Cohen’s *d* = 0.827)). There was no significant main effect of Stimulation Group or any two- or three-way interaction (all *p* > 0.1). Despite the lack of main effect of Stimulation Group, pairwise comparisons were still conducted in order to reassure that magnitudes were matched across stimulation groups. No significant differences were found between any two groups in each ear and harmonic condition (all *p* > 0.1, FDR-corrected according to multiple number of comparisons in each condition (i.e., ten)).

**Figure 3.**
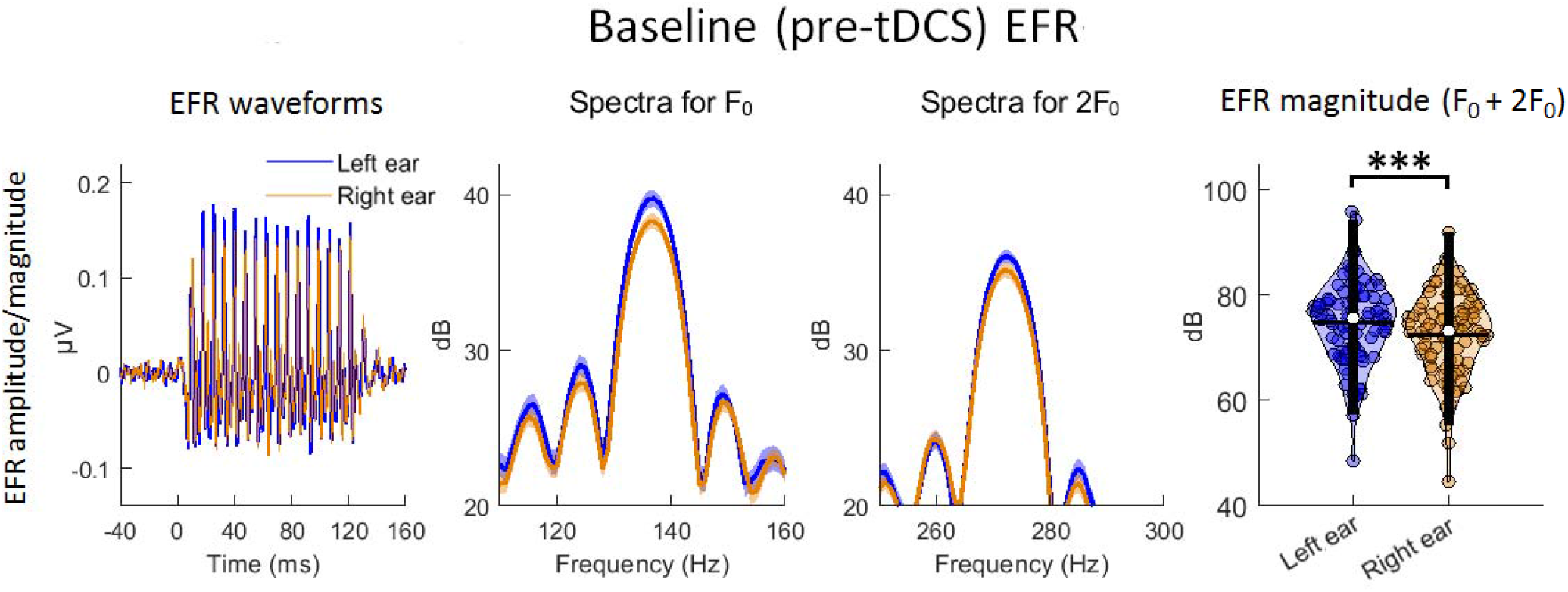
Comparison of baseline EFR magnitude between the left and the right ear conditions. From left to right shows the EFR waveforms, log-transformed FFT spectra for EFR at F_0_ and 2F_0_, and the data distributions of EFR magnitudes. The EFR waveforms were the temporal grand-averaged waveforms across all participants. FFT spectra were obtained based on the windowed EFR (zero-padded to 1 second) time-locked to the optimal neural lags (for F_0_ and 2F_0_, respectively; see the main text). The spectra show magnitudes peaking at 136 Hz (for F_0_) and 272 Hz (for 2F_0_), respectively. Data distributions were illustrated as violin plots based on collapsing the two harmonics (addition of EFR magnitudes at F_0_ and 2F_0_) for each participant due to the lack of significant interaction involving Harmonic. In each violin plot, the white dot in the middle refer to the median value; the horizontal and vertical lines indicate the ±1.5 interquartile range and the mean value, respectively. *Blue* and *brown* lines/dots denote data in the left and the right ear conditions, respectively. Shaded areas in the spectra cover the ranges of ±1 standard error (SE) from the means. \*\*\**p* < 0.001.

For the EFR neural lag, no significant main effects nor any interactions were found (all *p* > 1). Similar to EFR magnitude, pairwise comparisons were still conducted for the neural lag despite the lack of main effect of Stimulation Group. No significant differences were found between any two groups in each ear and harmonic condition (all *p* > 0.1, FDR corrected).

### After-effects across stimulation groups and ears

**Figure 4** shows the waveforms and FFT-power spectra of EFR pre- and post-tDCS in both ears, visualizing the after-effects across stimulation groups and ears. Linear mixed-effect regressions were conducted for the z-normalized tDCS after-effects on EFR. Statistical results are summarized in **Table 1**.

**Figure 4.**
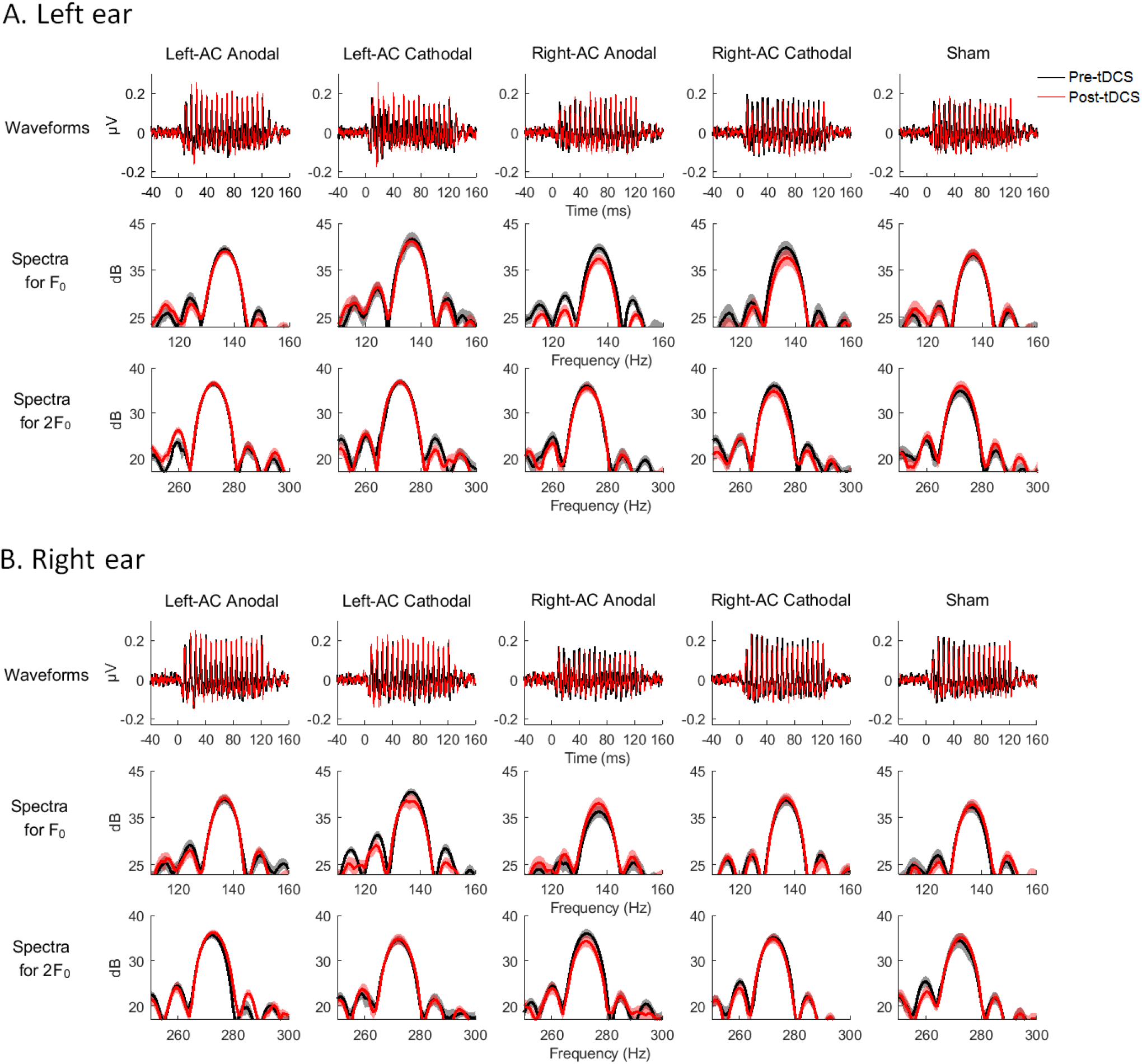
Pre- and post-tDCS waveforms and power spectra for EFR. **(A)** and **(B)** show the waveforms and log-transformed FFT-power spectra in the left and right ear condition, respectively. *Black* and *red* lines indicate the pre- and post-tDCS data, respectively. From left to right shows different stimulation groups (Left-AC Anodal, Left-AC Cathodal, Right-AC Anodal, Right-AC Cathodal and Sham). *Upper* panels show the grand-averaged EFR waveforms. *Mid* and *lower* panels show the FFT spectra based on the windowed EFR time-locked to the optimal neural lags for F_0_ (*mid*) and 2F_0_ (l*ower*). Shaded areas in the spectra cover the ranges of ±1 SE from the means.

**Table 1.** Statistics results for linear mixed-effect regressions for the after-effects (post-vs. pre-tDCS, z-normalized) on EFR magnitude and neural lag. . Stimulation Group, Ear and Harmonic were the fixed-effect factors and Participant was the random-effect factor. Significant *p* values (< 0.05) are in bold. \**p* < 0.05, \*\**p* < 0.01, \*\*\**p* < 0.001.

| DV | Fixed-effect factors/interactions | df1 | df2 | F | $p$ |
| --- | --- | --- | --- | --- | --- |
| After-effect on EFR magnitude (z-normalized) | Stimulation Group × Ear | 4 | 238.131 | 3.045 | **0.018\*** |
| | Stimulation Group × Harmonic | 4 | 238.131 | 0.472 | 0.756 |
| | Ear × Harmonic | 1 | 238.131 | 0.403 | 0.526 |
| | Stimulation Group × Ear × Harmonic | 4 | 238.131 | 1.541 | 0.191 |
|  | Stimulation Group | 4 | 136.312 | 3.503 | **0.009\*\*** |
|  | Ear | 1 | 238.131 | 16.037 | **$< 10^{-4}$\*\*\*** |
|  | Harmonic | 1 | 238.131 | 1.499 | 0.222 |
| After-effect on<br>EFR neural lag<br>(z-normalized) | Stimulation Group × Ear | 4 | 285.687 | 0.709 | 0.586 |
| | Stimulation Group × Harmonic | 4 | 129.520 | 1.080 | 0.369 |
| | Ear × Harmonic | 1 | 295.884 | 7.887 | **0.005\*\*** |
| | Stimulation Group × Ear × Harmonic | 4 | 295.884 | 1.016 | 0.399 |
|  | Stimulation Group | 4 | 140.489 | 0.861 | 0.489 |
|  | Ear | 1 | 285.687 | 0.156 | 0.693 |
|  | Harmonic | 1 | 129.520 | 1.397 | 0.239 |

For the after-effects on EFR magnitude, there were significant main effects of Stimulation Group (F(4, 136.312) = 3.503, *p* = 0.009) and Ear (F(1, 238.131) = 16.037, *p* <10^-4^) and a significant [Stimulation Group × Ear] interaction (F(1, 238.131) = 3.045, *p* = 0.018). Post-hoc analyses following the significant interaction were conducted via splitting the dataset by Ear. Specifically, additional linear mixed-effect regressions were conducted in the left ear and right ear conditions, respectively. A significant main effect of Stimulation Group was found in the left (F(4, 85) = 3.627, *p* = 0.009) but not the right ear condition (F(4, 123.058) = 0.919, *p* = 0.455). Pairwise comparisons were thus conducted between different stimulation groups using independent-sample t-tests in the left ear condition (collapsing the two harmonics due to no significant interactions involved with Harmonic). We found that Right-AC Anodal and Right-AC Cathodal resulted in significant decreases in EFR magnitude compared to Sham with large effect sizes (Right-AC Anodal vs. Sham: t(34) = -3.100, *p* = 0.019, Cohen’s *d* = 1.033; Right-AC Cathodal vs. Sham: t(34) = -3.351, *p* = 0.019, Cohen’s *d* = 1.117; *p* values were FDR corrected according to multiple number of comparisons (i.e., ten)). In the right ear condition, pairwise comparisons between stimulation groups were also conducted despite insignificant main effect of Stimulation Group to further reassure that after-effects did not differed between groups. Indeed, no significant differences were found between any two groups (all *p* > 0.5, FDR corrected). Data distributions of the after-effects on EFR magnitudes in the respectively ear conditions are illustrated as **Figure 5**.

**Figure 5.**
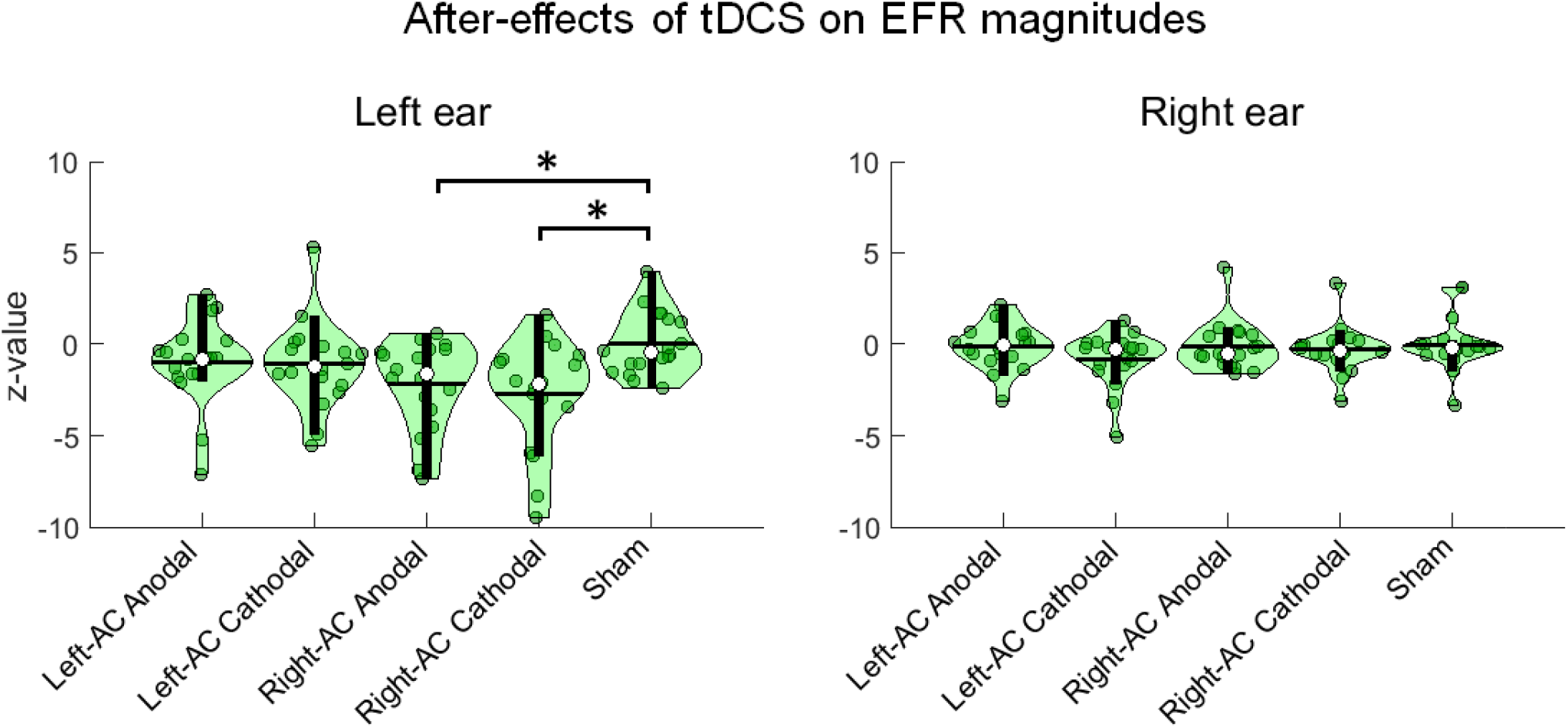
Data distributions of tDCS after-effects on EFR magnitudes across stimulation groups. After-effects are shown as z-values normalized according to Sham in the respective ear conditions. The distributions are illustrated based on collapsing the two harmonics due to the lack of significant interaction involving Harmonic. Data of different ear conditions are shown due to the significant [Stimulation Group × Ear] interaction. Post-hoc analyses following the interaction showed a significant main effect of Stimulation Group in the left but not the right ear condition. Pairwise comparisons showed significant decreases in EFR magnitudes for the Right-AC Anodal and Right-AC Cathodal compared to Sham in the left ear condition. \**p* < 0.05, FDR-corrected according to multiple number of comparisons in each ear condition (i.e., ten).

For the after-effects on EFR neural lag, no significant main effects nor interactions were found, except for the [Ear × Harmonic] interaction (F(1, 295.884) = 7.887, *p* = 0.005). Post-hoc analyses were thus required via splitting the dataset by Ear and Harmonic. Such effect, however, did not involve differences between stimulation groups and thus were not within the interest of the current study. The post-hoc results are therefore not reported here. In addition, to further reassure that after-effects on neural lag did not differ significantly between stimulation groups, pairwise comparisons were still conducted in the respective ear conditions despite no significant main effects/interactions related to Stimulation Group were found. No significant differences were found between any stimulation groups in either ear (all *p* > 0.4, FDR corrected).

Furthermore, the same linear mixed-effect regressions were replicated by further including potential confounding factors (Age, PTA (pre-tDCS), change in PTA (post- vs pre-tDCS) and HI) as fixed-effect covariates (including one confounder at a time). None of these factors changed the statistical significance of the original results and no significant main effects of them were found (all *p* > 0.2).

### Laterality and contralaterality of after-effects

Linear mixed-effect regressions were conducted for the after-effects on EFR magnitude with data of the Sham group excluded. As mentioned in *Materials and Methods*, neither Stimulation Type nor Harmonics had main effects or interacted significantly with other factors. To reduce analysis complexity, data of Anodal and Cathodal were grouped together and the two harmonics were collapsed before the regression analysis. Statistical results are shown in **Table 2**.

**Table 2.** Statistics results for linear mixed-effect regressions for the after-effects (post-vs. pre-tDCS, z-normalized) on EFR magnitude excluding the Sham group. . Stimulated Hemisphere and Contralaterality were the fixed-effect factors and Participant was the random-effect factor. Significant *p* values (< 0.05) are in bold. \**p* < 0.05, \*\*\**p* < 0.001.

| DV | Fixed-effect factors/interactions | df1 | df2 | F | $p$ |
| --- | --- | --- | --- | --- | --- |
| After-effect on EFR magnitude (z-normalized) | Stimulated Hemisphere × Contralaterality | 1 | 70 | 18.394 | **$< 10^{-4}***$** |
|  | Stimulated Hemisphere | 1 | 70 | 2.863 | 0.095 |
|  | Contralaterality | 1 | 70 | 6.456 | **0.013\*** |

A significant main effect of Contralaterality (F(1, 70) = 6.456, *p* = 0.013) and [Stimulated Hemisphere × Contralaterality] interaction (F(1, 70) = 18.394, *p* < 10^-4^) were found. Post-hoc pair-wise comparisons were conducted following the significant interaction comparing: (1) after-effects between different stimulated hemispheres (Left-AC vs Right-AC tDCS) in the ipsilateral and contralateral ear conditions, respectively, and (2) after-effects between ipsilateral and contralateral ear conditions for Left-AC and Right-AC tDCS, respectively. For (1), there was a significantly greater decrease in EFR magnitude for the Right-AC tDCS than the Left-AC tDCS in the contralateral ear condition (t(52.421) = -3.948, *p* < 0.001, unequal variance due to statistical significance of Levene’s test; Cohen’s *d* = 0.913), but not the ipsilateral ear. For (2), there was a significantly greater decrease in EFR magnitude in the contralateral than the ipsilateral ear for the Right-AC tDCS (t(50.456) = -4.514, *p* < 10^-4^, unequal variance due to statistical significance of Levene’s test; Cohen’s *d* = 1.064), but not Left-AC tDCS. Note that both significant effects had large effect sizes (Cohen’s *d* > 0.8). Data distributions of the after-effects across stimulated hemispheres and the ipsilateral and contralateral ear conditions are illustrated as **Figure 6**.

**Figure 6.**
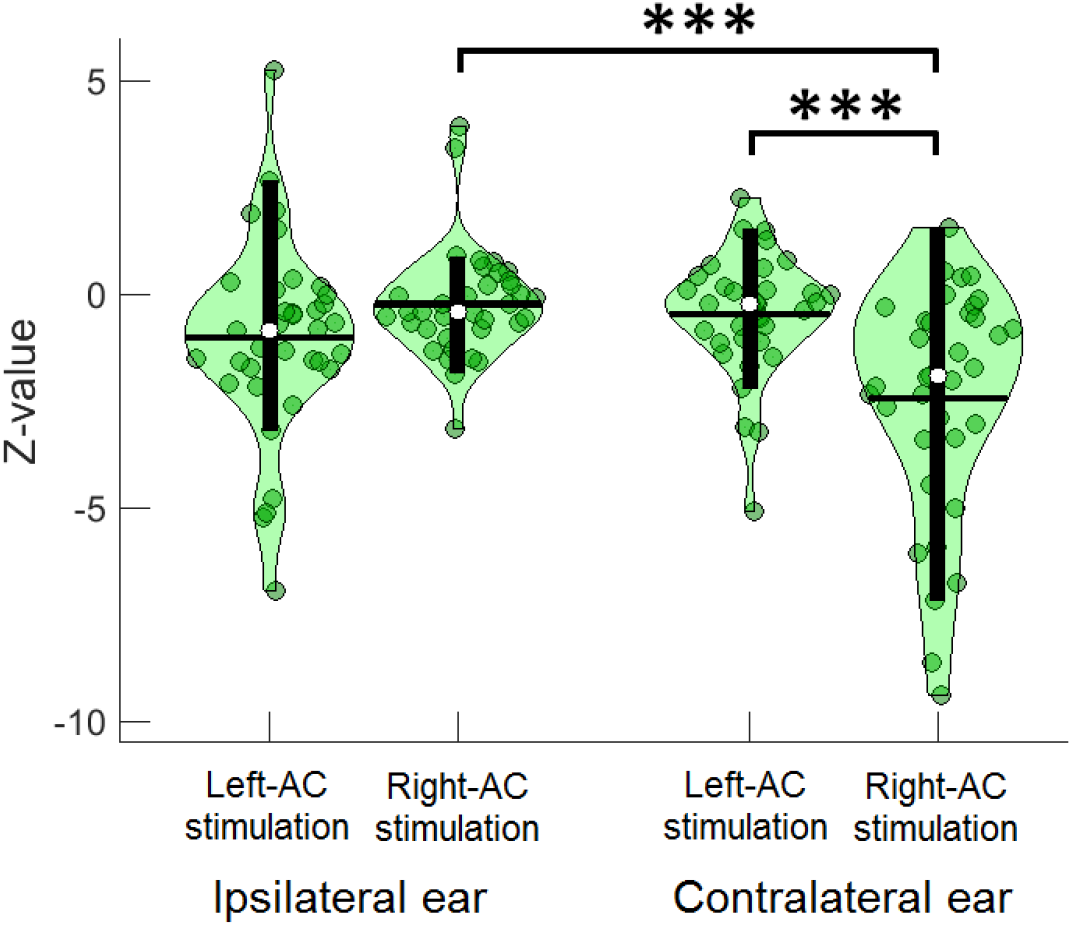
Data distributions of tDCS after-effects on EFR magnitudes with data of Sham excluded. After-effects are shown as z-values split by Stimulated Hemisphere (left/right auditory cortex), Contralaterality (ears ipsilateral/contralateral to the stimulated hemispheres). As neither Stimulation Type (Anodal/Cathodal) nor Harmonics (F_0_/2F_0_) had main effects or interacted with other factors, data of Anodal and Cathodal were group together and the two harmonics were collapsed. A significant [Stimulated Hemisphere × Contralaterality] was found. Post-hoc analyses following the interaction showed a significant greater decrease in the EFR magnitude for tDCS over the right (Right-AC tDCS) than the left auditory cortex (Left- AC tDCS) in the contralateral, but not ipsilateral ear condition; and a significant greater decrease in magnitude for the Right-AC than the Left-AC tDCS in the contralateral than the ipsilateral ear. \*\*\**p* < 0.001.

### Timing characteristics of significant after-effects

Independent sample t-tests were conducted in different time frames comparing z-values between the Right-AC tDCS and Sham in the left ear condition (**Figure 7A**), and between the Right-AC and Left-AC tDCS in their respective contralateral ear conditions (**Figure 7B**). The comparisons were conducted when combining the two harmonics (F_0_ plus 2F_0_) (see upper panels of **Figure 7**) and for F_0_ only (see lower panels of **Figure 7**). The results show that significant after-effects started to emerge from Frame 2 (7.4–14.8 ms after stimulus onset) for all comparisons (also see **Table 3** for the t-statistics). *P* values for the t-tests were FDR-corrected according to the total number of frames (i.e., seven). All significant effects had medium to large effect sizes (all Cohen’s *d* were in the range of either 0.5–0.8 or > 0.8, see **Table 3**). The results thus indicate that emergence of significant after-effects started at a relatively early stage (before 15 ms after stimulus onset, which is likely at the subcortical level, see Chandrasekaran and Kraus, 2010) along the central auditory systems.

**Figure 7.**
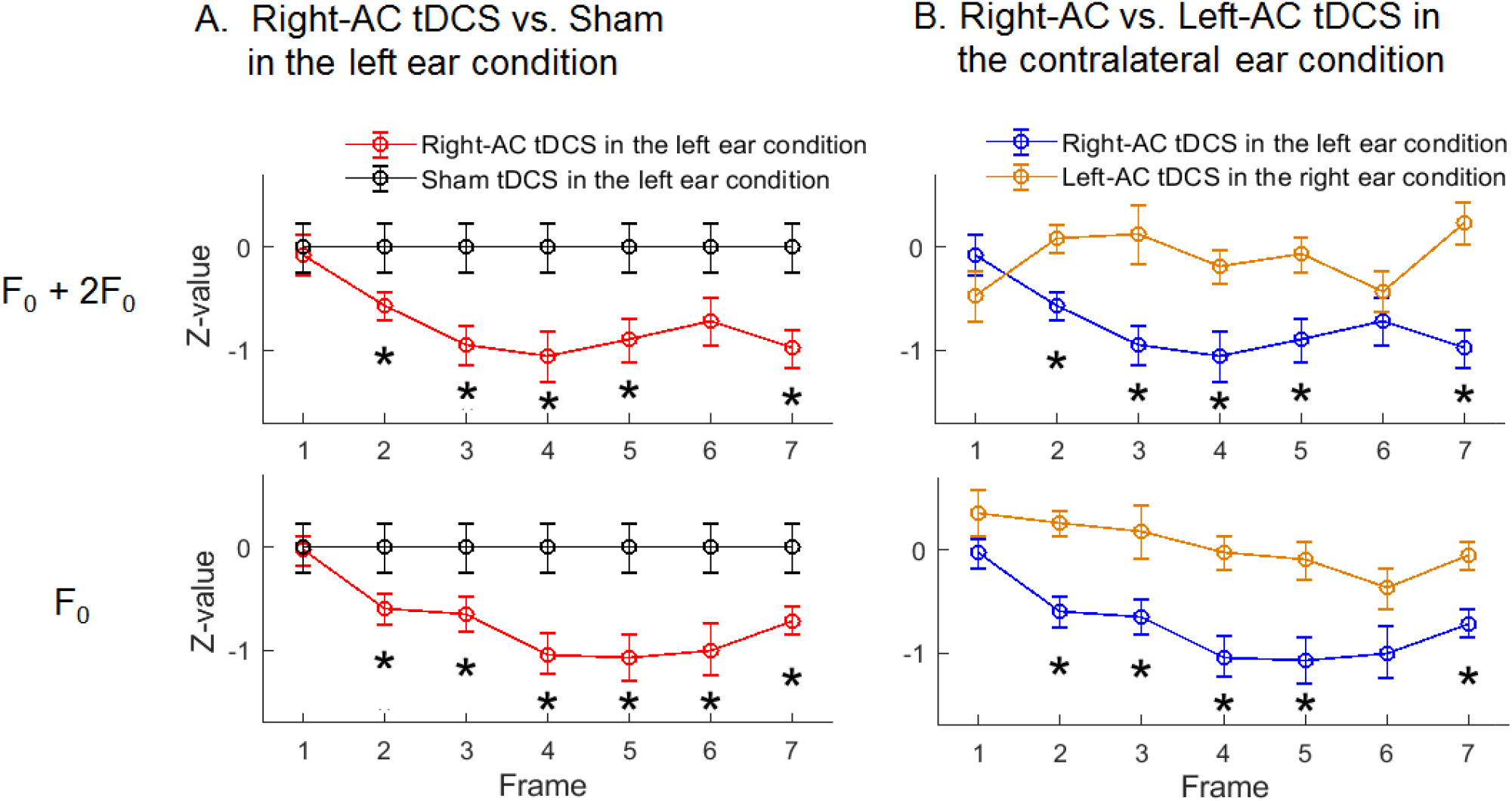
Timing characteristics of the after-effects and hemispheric laterality. Z-values are shown for the magnitudes of EFR amplitude profile that compare: **(A)** Right-AC tDCS with Sham in the left ear condition and **(B)** Right-AC with Left-AC tDCS in the respective contralateral ear conditions across time frames (Frame 1 to 7 time-locked to stimulus onset). The length of each frame was 7.4 ms corresponding to one cycle of F_0_. *Upper* and *lower* panels respectively indicate the comparisons when collapsing the magnitudes at the two harmonics (F_0_ + 2F_0_) and at F_0_ only. Error bars indicate the standard errors. Asterisks indicate the significant differences for each frame via independent-sample t-tests. Note that the mean values of Sham were all zeros because they were z-normalized in each frame. \**p* < 0.05, FDR corrected according to multiple number of comparisons (i.e., seven).

**Table 3.** Comparisons of after-effects on EFR magnitudes across time. Comparisons were conducted between Right-AC tDCS and Sham in the left ear condition (Comparison (a)) and between Right-AC tDCS and Left-AC tDCS in their respective contralateral ear condition (Comparison (b)). Independent t-tests were conducted from Frame 1 to Frame 7 that combined the two harmonics (F_0_ + 2F_0_) and at F_0_ only. The statistical results in the table correspond to the significant results illustrated in **Figure 7** (Comparison (a) corresponds to **Figure 7A** and Comparison (b) corresponds to **Figure 7B**). Numbers in the brackets denote the range for each time frame after the stimulus onset. The length of each frame was 7.4 ms (one cycle of F_0_). T, *p* and *d* refer to the t values, *p* values, and Cohen’s *d*, respectively. All *p* values were FDR corrected according to the number of comparisons (i.e., eight). Significant *p* values (< 0.05) are in bold. \**p* < 0.05, \*\**p* < 0.01, \*\*\**p* < 0.001.

|  |  |  | Frame 1 | Frame 2 | Frame 3 | Frame 4 | Frame 5 | Frame 6 | Frame 7 |
| --- | --- | --- | --- | --- | --- | --- | --- | --- | --- |
|  |  |  | (0–7.4 ms) | (7.4–14.8 ms) | (14.8–22.2 ms) | (22.2–29.6 ms) | (29.6–37 ms) | (37–44.4 ms) | (44.4–51.8 ms) |
| Comparison | F<sub>0</sub>+2F<sub>0</sub> | T | -0.236 | -2.267 | -2.948 | -2.796 | -2.652 | -1.929 | -3.200 |
| (a) |  | *p* | 0.815 | **0.028\*** | **0.005\*\*** | **0.007\*\*** | **0.011\*** | 0.059 | **0.002\*\*** |
|  |  | *d* | 0.068 | 0.654 | 0.851 | 0.807 | 0.766 | 0.557 | 0.924 |
|  | F<sub>0</sub> | T | -0.121 | -2.146 | -2.173 | -3.188 | -2.988 | -2.518 | -2.760 |
|  |  | *p* | 0.904 | **0.037\*** | **0.034\*** | **0.002\*\*** | **0.004\*\*** | **0.015\*** | **0.008\*\*** |
|  |  | *d* | 0.035 | 0.620 | 0.627 | 0.920 | 0.863 | 0.727 | 0.797 |
| Comparison | F<sub>0</sub>+2F<sub>0</sub> | T | 1.248 | -3.460 | -3.114 | -2.993 | -3.084 | -0.930 | -4.412 |
| (b) |  | *p* | 0.216 | **<0.001\*\*\*** | **0.003\*\*** | **0.004\*\*** | **0.003\*\*** | 0.355 | **<10<sup>-4</sup>\*\*\*** |
|  |  | *d* | 0.294 | 0.816 | 0.734 | 0.705 | 0.727 | 0.219 | 1.040 |
|  | F<sub>0</sub> | T | -1.464 | -4.279 | -2.689 | -3.910 | -3.347 | -1.974 | -3.352 |
|  |  | *p* | 0.148 | **<10<sup>-4</sup>\*\*\*** | **0.009\*\*** | **<0.001\*\*\*** | **0.001\*\*** | 0.052 | **0.001\*\*** |
|  |  | *d* | 0.345 | 1.009 | 0.634 | 0.922 | 0.789 | 0.465 | 0.790 |

### Pitch discrimination

A linear mixed-effect regression was conducted for the change in pitch discrimination performances. The results showed no significant main effect of Stimulation Group (F(4, 85) = 0.393) and no significant differences in the change in performances between any two stimulation groups (all *p* > 0.4, FDR corrected). Also Pearson’s correlations were conducted between the changes in performances and the after-effects on EFR magnitude (collapsing the two harmonics) in both ears in the respective stimulation groups. No significant correlations were found (all *p* > 0.3, FDR corrected).

### Summary of results

To summarize, our results showed as follows. First, we confirmed that EFR magnitude and neural lag were matched across stimulation groups at the baseline, so any after-effect should not be attributed to baseline differences. We further found a relatively small (small to medium effect size) but significant left ear laterality for the baseline EFR magnitude. Second, we showed a significant [Stimulation Group × Ear] interaction for the after-effects on EFR magnitude. Post-hoc analyses found that tDCS over the right auditory cortex resulted in significant changes (decreases) in magnitude compared to Sham in the left ear (contralateral to the right auditory cortex) condition. On the other hand, no significant after-effects were found for tDCS over the left auditory cortex compared to Sham. Third, we showed a significant [Stimulated Hemisphere × Contralateral] interaction and a hemispheric laterality in which the decrease in EFR magnitude in the contralateral ear was significantly greater when tDCS was applied over the right than the left auditory cortex. Finally, analyses on timing characteristics showed that emergence of the significant after-effects and laterality mentioned in the second and the third points started from the relatively early, likely subcortical stage in the auditory systems.

The results therefore addressed our hypotheses that tDCS over the right auditory cortex causes changes in EFR along the contralateral auditory. It thus validates the causal relationship between the right auditory cortex and EFR.

## Discussion

The current study used a combined tDCS and EEG approach to test for a causal contribution of auditory cortex to speech-evoked EFR in healthy right-handed participants. This approach can inform us that the cortical contribution found in the previous studies is not merely a result of specific source localization techniques or observed association between cortical activations and EFR, but that EFR is casually related to neuroplastic changes induced by neurostimulation in the auditory cortex. The left and right auditory cortices were neurostimulated in different groups of participants and the after-effects of tDCS on the EFR were examined during monaural listening to a repeated speech syllable. Our results showed that tDCS over the right auditory cortex resulted in significant decrease in EFR magnitude compared to Sham as well as tDCS over the left auditory cortex (i.e., hemispheric laterality) when the listening ear was contralateral to the stimulated site. No significant after-effects were found for tDCS over the left auditory cortex. The results thus agree with previous studies that have shown a close relationship between the right auditory cortex and EFR (Coffey et al., 2016, 2017a, 2017b; Hartmann and Weisz, 2019) and provide the very first evidence for a causal relationship.

### Causal contributions of the right auditory cortex to EFR and the hemispheric laterality

A recent effort was made to study how EFR is modulated by inhibiting excitability of the right auditory cortex via repetitive transcranial magnetic stimulation (rTMS) (Lopez-Caballero et al., 2020). rTMS was applied over the right auditory cortex and EFR magnitudes obtained via EEG were compared between post- and pre-TMS while participants binaurally listened to repeated syllables. No significant after-effects on EFR were found. Here, we argue that the right auditory cortex may contribute to EFR mainly along the contralateral pathway. If this is the case, the insignificant effect from the ipsilateral ear during binaural listening could conceal the genuine impact of the neuro-stimulation. We also argue that it is more appropriate to also include the condition where stimulation is applied over the left auditory cortex to test whether the after-effect occurs specifically in the right hemisphere and to test for hemispheric laterality. The present study, therefore, used an approach in which participants monaurally listened to stimuli to allow for testing contralaterality and neuro-stimulations were applied over both left and right auditory cortices so that hemispheric laterality were directly studied.

We found that tDCS over the right, but not left, auditory cortex resulted in significant changes in EFR magnitude compared to Sham in its contralateral (i.e., left) ear condition. Together with the hemispheric laterality found in the contralateral ear condition, our results argue for a causal role of the right auditory cortex in processing speech periodicity along the contralateral pathway in the central auditory systems. We suggest that the present study advances our understanding of the relationship between EFR and pitch processing in the right auditory cortex. Previous studies have shown that EFR is closely related to pitch perception. EFR strength can be enhanced by both short-term perceptual training of pitch discrimination (Carcagno and Plack, 2011) as well as long-term musical experience (Musacchia et al., 2007; Wong et al., 2007; Strait et al., 2009; Bidelman et al., 2011). Furthermore, EFR has been used as an index of neural fidelity of linguistic pitch and the fidelity is greater in tonal language than in non-tonal language speakers (Krishnan et al., 2004, 2005, 2009). Despite this, however, rather than reflecting the result of pitch extraction, EFR has been suggested to reflect subcortical responses to monaural temporal information (e.g., periodicity cues) that are important for extracting pitch of complex sounds (i.e., ‘pitch-bearing’ information; Gockel et al., 2011). On the other hand, the process of pitch extraction itself takes place in the auditory cortex (Penagos et al., 2004; Bendor and Wang, 2005; Puschmann et al., 2010) with a right hemispheric specialization (Zatorre and Berlin, 2001; Patterson et al., 2002; Hyde et al., 2008; Mathys et al., 2010; Albouy et al., 2013; Matsushita et al., 2015, 2021). In this respect, the current after-effects of tDCS may reflect a top-down corticofugal modulation process in which the right auditory cortex affects the processing of pitch-bearing information that occurs at the subcortical level. A model of top-down modulation between auditory cortex and subcortex that controls the temporal dynamics of pitch processing has been proposed and stresses that this process is key for pitch perception (Balaguer-Ballester et al., 2006). Alternatively, although EEG mainly captures EFR signals originating from the brainstem (Bidelman, 2015, 2018), cortical sources have been found dominated in the right hemisphere (Coffey et al., 2016; 2017a). It may be that the after-effects reflected the changes in the cortical EFR activities. Therefore, additional analyses for the timing characteristics of when significant after-effects started to emerge were further investigated in the current study. The results seem to support the proposal arguing for the top-down role of the auditory cortex. This will be further discussed in the next section (*Timing characteristics of the significant after-effects*).

It is also noteworthy that we showed a laterality effect at the baseline as well, where EFR had significantly greater magnitude in the left than the right ear condition. This echoes previous findings showing right-hemispheric laterality for auditory steady-state responses (ASSRs) (but at lower frequencies of 40 and 80 Hz, Ross et al., 2005; Luke et al., 2017; Vanvooren et al., 2014). Both ASSRs and EFR are phase-locked responses to envelope modulations, implying that phase-locked envelope responses in general might be more prominent along the contralateral pathway from the left ear to the right auditory cortex. However, it is not clear whether and how auditory cortex contributes to this laterality for EFR. As such, the current results provide confirmatory evidence for a causal cortical contribution to this laterality. It would be interesting to see whether such causality could be replicated for ASSR in the future. It should also be noted that the present study was conducted specifically in right-handed participants. While hemispheric laterality is associated with handedness (Carey and Johnston, 2014; Willems et al., 2014), the current results may not necessarily apply to people who are left-handed or ambidextrous.

Besides measuring EFR, the present study also used a pitch discrimination task during tDCS application so that the auditory cortex was kept active and the tDCS effects were optimized (Reato et al., 2010; Ranieri et al., 2012; Bikson and Rahman, 2013). Unlike EFR, changes in pitch discrimination performances did not differ between stimulation groups and no correlations were found between changes in performances and EFR magnitudes in any group. This may be because, with our electrode configuration (active electrode on the auditory cortex and reference on the forehead above the contralateral eyebrow), currents generated by tDCS would pass through not only auditory cortices but also various brain regions due to its marked diffusive nature (Faria et al., 2011; Bai et al., 2014; Unal and Bikson, 2018). Pitch discrimination involves not only sensory regions (i.e., auditory cortices) but also higher-order regions such as frontal and prefrontal cortices (Zatorre et al., 1992; Palomar-Garcia, 2020) and anterior cingulate cortex (Schwenzer and Mathiak, 2011) which may have been stimulated by tDCS. We thus argue that it is not surprising that no significant results were found for pitch discrimination because of such diffusive effects of tDCS.

Another important issue needs discussion is that the present study focused on EFR that reflects neural encoding of periodicity envelope at F_0_ and its harmonic (2F_0_). Besides periodicity envelope, temporal fine structure (TFS), particularly at the resolved harmonic region, is also essential for pitch perception (Moore et al., 2014). Here, we did not include responses to TFS because they mainly reflects encoding of sounds at the auditory periphery or cochlear microphonics (Aiken and Picton, 2008) rather than responses at the central auditory systems. Also, there may be electrical artefacts when recording responses to TFS, while EFR was obtained by adding responses to the positive and negative polarities such that artefacts are minimized (Skoe and Kraus, 2010). It is therefore valuable for the future to explore an artefact-free response to TFS located in the central auditory systems and how it may relate to cortical hemispheric laterality. In contrast to TFS, periodicity envelope is most important for pitch perception when auditory signals are dominated by unresolved harmonics (Oxenham et al., 2009; Moore and Gockel, 2011). Furthermore, periodicity encoding is more important when TFS is not well perceived or available in clinical populations. For example, evidence showed that encoding of periodicity envelope is less deteriorated than TFS by ageing and hearing loss (Halliday et al., 2019, Moore, 2021). In hearing protheses such as cochlear implants (CI), periodicity envelope rather than TFS is effectively conveyed, so CI recipients would rely mainly on periodicity encoding to perceive pitch (Hofmann and Wouters, 2010; Wouters et al., 2015; Gransier et al., 2020). Therefore, it would be also meaningful for future studies to investigate cortical contributions to EFR in hearing-impaired populations.

### Timing characteristics of the significant after-effects

Analyses of the timing characteristics showed that the significant after-effects and hemispheric laterality observed in the present study emerged at a relative early, subcortical time window (∼7-15 ms after stimulus onset). This early emergence happened both when collapsing EFR magnitudes at F_0_ and 2F_0_ and for magnitudes at F_0_ only.

Therefore, such results indicate that the after-effects may be introduced via a top-down corticofugal modulation process in which excitability changes in the right auditory cortex affected EFR that occurs at the subcortical level. It is noteworthy that ‘cortical contribution’ we refer to here is not limited to possible contributions of cortical sources of EFR, but also corticofugal modulation that involves changes in the right auditory cortex. Another argument may be that, although tDCS was applied over the auditory cortices, the after-effects could be due to the impact of tDCS *directly* on the brainstem without cortical contributions. However, we argue that this latter proposal was not likely to be the case, because if it were, impacts imposed by tDCS on the brainstem are expected to be similar between stimulations over different hemispheres. In such case, the laterality of after-effects should be determined only by factors below the cortical level (i.e., ear conditions) regardless of which hemispheres were stimulated. Our current results, however, showed a significant interaction between Stimulated Hemisphere and Contralaterality (ears contralateral to the stimulated hemisphere). Hemispheric laterality was shown in which tDCS resulted in significantly greater after-effects over the right than the left auditory cortex in the contralateral, but not ipsilateral, ear condition. This therefore argues against the possibility that the after-effects were consequences of the tDCS directly imposed on the brainstem and argues for the top-down corticofugal modulation process.

The auditory corticofugal modulation has been well established in animal studies (Suga et al., 2000; Bajo et al., 2010; Bajo and King, 2013; Suga, 2020). Specifically, excitatory electrical stimulations in the auditory cortex can facilitate the responses of inferior colliculus (IC) neurons that have the same tuning frequencies as the stimulated cortical neurons (Bajo and King, 2013; Suga, 2020). On the other hand, inactivating auditory cortex reduces the responses of IC neurons (Zhang and Suga, 1997; Yan and Suga, 1999; Bajo and King, 2013). Therefore, a possible mechanism that underlies the current observed after-effects would be that tDCS altered the excitability in the auditory cortex. This would then lead to hyperpolarization of subcortical postsynaptic neurons through the efferent connections from the auditory cortex to the subcortex. The hyperpolarization would raise the thresholds for the occurrence of action potentials in response to speech periodicity, reflected by the decrease in EFR magnitude. The interesting result we observed here is that this process happens specifically for the right auditory cortex along the contralateral pathway. The human auditory cortex has been shown to have sharper frequency tuning in the right than the left hemisphere (Liégeois-Chau et al., 1999; 2001), suggesting greater number of neurons responsible for spectral analyses of periodicity information that support pitch perception in the right hemisphere (Zatorre et al., 2002). Therefore, top-down corticofugal modulation for pitch perception (Balaguer-Ballester et al., 2006) may be prominent in the right hemisphere and hence more susceptible to external neuro- stimulations compared to Sham and stimulations over the left hemisphere.

Despite our observations of the timing characteristics, some limitations are yet to be addressed. First, previous research argued that cortical EFRs have upper frequency limit at ∼100 Hz (Bidelman, 2018). However, recent data have shown that cortical EFRs also include frequencies that cover a more common range of human vocal pitch (100-140 Hz, Ross et al., 2020; 100-200 Hz, Guo et al., 2020). The present study used a speech syllable with F_0_ at 136 Hz that falls within the 100-200 Hz range. Nonetheless, we had not obtained compelling results here in order to conclude that the after-effects were directly from the cortex. Future studies may use a lower F_0_ to clarify whether the timing characteristics are different across F_0_ frequencies. Second, using scalp-recorded EEG, we were not able to spatially localize the after- effects. Therefore, an even better approach for future studies to consider is to use technique like MEG that is capable of localizing the after-effects at the subcortical and/or cortical level (Coffey et al., 2016). Combining the timing and spatial information could provide a more comprehensive image showing us both when and where the effects start to emerge.

### Neurophysiological consequences of tDCS

An intriguing finding of the present study is that anodal and cathodal tDCS resulted in the same direction of changes, both causing decreases in EFR magnitude for tDCS over the right auditory cortex in the contralateral (i.e., left) ear condition. Conventionally, anodal and cathodal stimulations reflect depolarization and hyperpolarization of neurons, respectively, which should lead to opposite directions of after-effects (Jacobson et al., 2012). However, it is not unusual that tDCS has polarity-independent effects due to the underlying complexity of its neurophysiological consequences. For example, several studies have shown that anodal and cathodal tDCS have the same effects on excitability of motor cortex (Antal et al., 2007), motor learning (de Xivry et al., 2011), cerebellar functions for working memory (Ferrucci et al., 2008) and visuomotor learning (Shah et al., 2013). The mechanisms underlying excitatory/inhibitory consequences of anodal and cathodal stimulations may be the changes in concentrations of relevant neurotransmitters. Stagg et al., (2009) found that anodal tDCS over the motor cortex causes decreases in gamma-amino butyric acid (GABA) concentration that lead to excitation, whilst cathodal tDCS also causes decreases in GABA but greater concurrent decreases in glutamate that lead to cortical inhibition. A recent study applied tDCS over the human auditory cortex and found that both anodal and cathodal stimulations resulted in increases in relative concentration of GABA to glutamate (Heimrath et al., 2020). This is consistent with the current results that both anodal and cathodal tDCS led to decreased EFR magnitude, indicating that both stimulation types may have introduced neural inhibition in the auditory cortex. It would be worthwhile for future studies to investigate how changes in concentrations of neurotransmitters by neuro-stimulation relate to changes in EFR, which can help us better understand the underlying mechanisms of cortical contributions to EFR.

### Speech-specific or domain-general

Our auditory stimulus used to obtain EFR was a speech syllable. Another important question is whether our results are speech-specific or could be generalized to non-speech stimuli. It has been shown that the right auditory cortical contributions are similar when using a speech syllable and a music note (Coffey et al., 2017a). However, speech and music notes share a great deal of physical properties (e.g., spectral complexity and harmonic structures) which may lead to such similarity. It is not clear whether using other stimuli such as complex tones, amplitude-modulated tones and iterated rippled noise (Krishnan et al., 2009; Ananthakrishnan and Krishnan, 2018) would lead to the same or different effects as observed here. A future direction is then to examine whether the causal contribution of the right auditory cortex is speech-specific or domain-general.

## Conclusion

We showed that tDCS over the right, but not left, auditory cortex resulted in significant changes in speech-evoked EFR magnitude compared to Sham in the contralateral (i.e., left) ear condition. Crucially, we also showed a hemispheric laterality in which the after-effect was greater for tDCS applied over the right than the left auditory cortex in the contralateral ear condition. Furthermore, we showed that the after-effects and the laterality emerged from the relatively early, subcortical stages, indicating a top-down corticofugal modulation of the right auditory cortex on the subcortex. The current results thus validate the previous findings that the right auditory cortex makes significant contributions to EFR (Coffey et al., 2016, 2017a, 2017b; Hartmann and Weisz, 2019) by establishing a causal relationship between the two. To our knowledge, this is the first evidence for this causality. Our findings should advance the understanding of how periodicity and pitch information are processed along the central auditory pathways in the human brain.

## Notes

We thank Dr Naheem Bashir for instructions on tDCS setups and Mr Andrew Clark for technical support for EEG recording. *Conflict of Interest*: None declared.

## References

Aiken SJ, Picton TW. 2008. Envelope and spectral frequency-following responses to vowel sounds. Hearing Research. 245:35–47.

Albouy P, Mattout J, Bouet R, Maby E, Sanchez G, Aguera PE, Daligault S, Delpuech C, Bertrand O, Caclin A, Tillmann B. 2013. Impaired pitch perception and memory in congenital amusia: the deficit starts in the auditory cortex. Brain. 136(5):1639–1661.

Ananthakrishnan S, Krishnan A. 2018. Human frequency following responses to iterated rippled noise with positive and negative gain: Differential sensitivity to waveform envelope and temporal fine-structure. Hearing Research. 367:113–123.

Ananthakrishnan S, Krishnan A, and Bartlett E. 2016. Human frequency following response: neural representation of envelope and temporal fine structure in listeners with normal hearing and sensorineural hearing loss. Ear and Hearing. 37(2):e91.

Anderson S, Parbery-Clark A, Yi HG, Kraus N. 2011. A neural basis of speech-in-noise perception in older adults. Ear and Hearing. 32(6):750.

Anderson S, Parbery-Clark A, White-Schwoch T, Kraus N. 2012. Aging affects neural precision of speech encoding. Journal of Neuroscience. 32(41):14156–14164.

Antal A, Terney D, Poreisz C, Paulus W. 2007. Towards unravelling task-related modulations of neuroplastic changes induced in the human motor cortex. European Journal of Neuroscience. 26(9):2687–2691.

Bai S, Dokos S, Ho KA, Loo C. 2014. A computational modelling study of transcranial direct current stimulation montages used in depression. NeuroImage. 87:332–344.

Bajo VM, King AJ. 2013. Cortical modulation of auditory processing in the midbrain. Frontiers in Neural Circuits. 6:114.

Bajo VM, Nodal FR, Moore DR, King AJ. 2010. The descending corticocollicular pathway mediates learning-induced auditory plasticity. Nature Neuroscience. 13:253–260.

Balaguer-Ballester E, Clark NR, Coath M, Krumbholz K, Denham SL. 2009. Understanding pitch perception as a hierarchical process with top-down modulation. PLoS Computational Biology. 5(3):e1000301.

Banai K, Abrams D, Kraus N. 2007. Sensory-based learning disability: Insights from brainstem processing of speech sounds. International Journal of Audiology. 46(9):524–532.

Bendor D, Wang X. 2005. The neuronal representation of pitch in primate auditory cortex. Nature. 436(7054):1161–1165.

Bidelman GM, Krishnan A, Gandour JT. 2011. Enhanced brainstem encoding predicts musicians’ perceptual advantages with pitch. European Journal of Neuroscience. 33(3):530–538.

Bidelman GM. 2015 Multichannel recordings of the human brainstem frequency-following response: scalp topography, source generators, and distinctions from the transient ABR. Hearing Research. 323:68–80.

Bidelman GM. 2018. Subcortical sources dominate the neuroelectric auditory frequency- following response to speech. NeuroImage. 175:56–69.

Bidelman GM, Lowther JE, Tak SH, Alain C. 2017. Mild cognitive impairment is characterized by deficient brainstem and cortical representations of speech. Journal of Neuroscience. 37(13):3610–3620.

Bikson M, Rahman A. 2013. Origins of specificity during tDCS: anatomical, activity-selective, and input-bias mechanisms. Frontiers in Human Neuroscience. 7:688.

Boroda E, Sponheim SR, Fiecas M, Lim KO, 2020. Transcranial direct current stimulation (tDCS) elicits stimulus-specific enhancement of cortical plasticity. NeuroImage. 211:116598.

Carcagno S, Plack CJ, 2011, Subcortical plasticity following perceptual learning in a pitch discrimination task. Journal of the Association for Research in Otolaryngology. 12(1):89–100.

Carey DP, Johnstone LT. 2014, Quantifying cerebral asymmetries for language in dextrals and adextrals with random-effects meta analysis. Frontiers in Psychology. 5:1128.

Cha K, Zatorre RJ, Schönwiesner M. 2016. Frequency selectivity of voxel-by-voxel functional connectivity in human auditory cortex. Cerebral Cortex. 26(1):211–224.

Chandrasekaran B, Kraus N. 2010. The scalp-recorded brainstem response to speech: Neural origins and plasticity. Psychophysiology. 47(2):236–246.

Coffey EB, Herholz SC, Chepesiuk AM, Baillet S, Zatorre RJ. 2016. Cortical contributions to the auditory frequency-following response revealed by MEG. Nature Communications. 7(1):1–11.

Coffey EB, Musacchia G, Zatorre RJ. 2017a. Cortical correlates of the auditory frequency- following and onset responses: EEG and fMRI evidence. Journal of Neuroscience. 37(4):830–838.

Coffey EB, Chepesiuk AM, Herholz SC, Baillet S, Zatorre RJ. 2017b. Neural correlates of early sound encoding and their relationship to speech-in-noise perception. Frontiers in Neuroscience. 11:479.

Coffey EB, Nicol T, White-Schwoch T, Chandrasekaran B, Krizman J, Skoe E, Zotorre RJ, Kraus N. 2019. Evolving perspectives on the sources of the frequency-following response. Nature Communications. 10(1):1–10.

Cunningham J, Nicol T, Zecker SG, Bradlow A, Kraus N. 2001. Neurobiologic responses to speech in noise in children with learning problems: deficits and strategies for improvement. Clinical Neurophysiology. 112(5):758–767.

de Xivry JJ, Marko MK, Pekny SE, Pastor D, Izawa J, Celnik P, Shadmehr R. 2011. Stimulation of the human motor cortex alters generalization patterns of motor learning. Journal of Neuroscience. 31(19):7102–7110.

Dimitrijevic A, John MS, Picton TW. 2004. Auditory steady-state responses and word recognition scores in normal-hearing and hearing-impaired adults. Ear and Hearing. 25(1):68–84.

Encina-Llamas G, Harte JM, Dau T, Shinn-Cunningham B, Epp B. 2019. Investigating the effect of cochlear synaptopathy on envelope following responses using a model of the auditory nerve. Journal of the Association for Research in Otolaryngology. 20(4):363–382.

Faria P, Hallett M, Miranda PC, 2011. A finite element analysis of the effect of electrode area and inter-electrode distance on the spatial distribution of the current density in tDCS. Journal of Neural Engineering. 8(6):066017.

Ferrucci R, Marceglia S, Vergari M, Cogiamanian F, Mrakic-Sposta S, Mameli F, Zago S, Babieri S, Priori A. 2008. Cerebellar transcranial direct current stimulation impairs the practice-dependent proficiency increase in working memory. Journal of Cognitive Neuroscience. 20(9):1687–1697.

Filmer HL, Dux PE, Mattingley JB. 2014. Applications of transcranial direct current stimulation for understanding brain function. Trends in Neurosciences. 37(12):742–753.

Fujihira H, Shiraishi K. 2015. Correlations between word intelligibility under reverberation and speech auditory brainstem responses in elderly listeners. Clinical Neurophysiology, 126(1):96–102.

Gockel HE, Carlyon RP, Mehta A, Plack CJ. 2011. The frequency following response (FFR) may reflect pitch-bearing information but is not a direct representation of pitch. Journal of the Association for Research in Otolaryngology. 12(6):767–782.

Gorina-Careta N, Kurkela JL, Hamalainen J, Astikainen P, Escera C. 2021. Neural generators of the frequency-following response elicited to stimuli of low and high frequency: A magnetoencephalographic (MEG) study. NeuroImage. 231:117866.

Gransier R, Carlyon RP, Wouters J. 2020. Electrophysiological assessment of temporal envelope processing in cochlear implant users. Scientific Reports. 10(1):1–14.

Guo N, Si X, Zhang Y, Ding Y, Zhou W, Zhang D, Hong B. 2021. Speech frequency- following response in human auditory cortex is more than a simple tracking. NeuroImage 226:117545.

Halliday LF, Rosen S, Tuomainen O, Calcus A. 2019. Impaired frequency selectivity and sensitivity to temporal fine structure, but not envelope cues, in children with mild-to- moderate sensorineural hearing loss. The Journal of the Acoustical Society of America. 146(6):4299–4314.

Hartmann T, Weisz N. 2019. Auditory cortical generators of the Frequency Following Response are modulated by intermodal attention. NeuroImage. 203:116185.

Heimrath K, Brechmann A, Blobel-Lüer R, Stadler J, Budinger E, Zaehle T. 2020. Transcranial direct current stimulation (tDCS) over the auditory cortex modulates GABA and glutamate: a 7 T MR-spectroscopy study. Scientific Reports. 10(1):1–8.

Herdman AT, Lins O, Van Roon P, Stapells DR, Scherg M, Picton TW. 2002. Intracerebral sources of human auditory steady-state responses. Brain Topography. 15(2):69–86.

Hofmann M, Wouters J. 2010. Electrically evoked auditory steady state responses in cochlear implant users. Journal of the Association for Research in Otolaryngology. 11(2):267–282.

Holmberg EB, Hillman RE, Perkell JS. 1988. Glottal airflow and transglottal air pressure measurements for male and female speakers in soft, normal, and loud voice. The Journal of the Acoustical Society of America. 84(2):511–529.

Hornickel J, Kraus N. 2013. Unstable representation of sound: a biological marker of dyslexia. Journal of Neuroscience. 33(8):3500–3504.

Hyde KL, Peretz I, Zatorre RJ. 2008. Evidence for the role of the right auditory cortex in fine pitch resolution. Neuropsychologia. 46(2):632–639.

Impey D, Knott V. 2015. Effect of transcranial direct current stimulation (tDCS) on MMN- indexed auditory discrimination: A pilot study. Journal of Neural Transmission. 122:1175–1185.

Jacobson L, Koslowsky M, Lavidor M. 2012. tDCS polarity effects in motor and cognitive domains: a meta-analytical review. Experimental Brain Research. 216(1):1–10.

Krishnan A, Xu Y, Gandour JT, Cariani PA. 2004. Human frequency-following response: representation of pitch contours in Chinese tones. Hearing Research. 189:1–12.

Krishnan A, Xu Y, Gandour J, & Cariani P. 2005. Encoding of pitch in the human brainstem is sensitive to language experience. Cognitive Brain Research. 25(1):161–168.

Krishnan A, Gandour JT, Bidelman GM, Swaminathan J. 2009. Experience dependent neural representation of dynamic pitch in the brainstem. Neuroreport. 20(4):408.

Kunze T, Hunold A, Haueisen J, Jirsa V, Spiegler A. 2016. Transcranial direct current stimulation changes resting state functional connectivity: a large-scale brain network modeling study. NeuroImage. 140:174–187.

Lehmann A., Schönwiesner M. 2014. Selective attention modulates human auditory brainstem responses: relative contributions of frequency and spatial cues. PloS One. 9(1):e85442.

Liegeois-Chauvel C, De Graaf JB, Laguitton V, Chauvel P. 1999. Specialization of left auditory cortex for speech perception in man depends on temporal coding. Cerebral Cortex. 9(5):484–96.

Liegeois-Chauvel C, Giraud K, Badier JM, Marquis P, Chauvel P. 2001. Intracerebral evoked potentials in pitch perception reveal a functional asymmetry of the human auditory cortex. Annals of the New York Academy of Sciences. 930(1):117–32.

Liu JG, Morgan GL. 2006. FFT selective and adaptive filtering for removal of systematic noise in ETM+ imageodesy images. IEEE Transactions on Geoscience and Remote Sensing. 44(12):3716–3724.

Lorenzi C, Gilbert G, Carn H, Garnier S, Moore BC. 2006. Speech perception problems of the hearing impaired reflect inability to use temporal fine structure. Proceedings of the National Academy of Sciences. 103(49):18866–18869.

Luke R, De Vos A, Wouters J. 2017. Source analysis of auditory steady-state responses in acoustic and electric hearing. NeuroImage. 147:568–576.

Mai G, Tuomainen J, Howell P. 2018. Relationship between speech-evoked neural responses and perception of speech in noise in older adults. The Journal of the Acoustical Society of America. 143(3):1333–1345.

Mai G, Schoof T, Howell P. 2019. Modulation of phase-locked neural responses to speech during different arousal states is age-dependent. NeuroImage. 189:734–744.

Mathys C, Loui P, Zheng X, Schlaug G. 2010. Non-invasive brain stimulation applied to Heschl’s gyrus modulates pitch discrimination. Frontiers in Psychology. 1:193.

Matsushita R, Andoh J, Zatorre RJ. 2015. Polarity-specific transcranial direct current stimulation disrupts auditory pitch learning. Frontiers in Neuroscience. 9:174.

Matsushita R, Puschmann S, Baillet S, Zatorre RJ. 2021. Inhibitory effect of tDCS on auditory evoked response: Simultaneous MEG-tDCS reveals causal role of right auditory cortex in pitch learning. NeuroImage. 233:117915.

Moore BC. 2014. Auditory processing of temporal fine structure: Effects of age and hearing loss. (World Scientific, Singapore, 2014).

Moore BC 2021. Effects of hearing loss and age on the binaural processing of temporal envelope and temporal fine structure information. Hearing Research. 402:107991.

Moore, BC, Gockel, HE. 2011. Resolvability of components in complex tones and implications for theories of pitch perception. Hearing Research. 276:88–97.

Musacchia G, Sams M, Skoe E, Kraus N. 2007. Musicians have enhanced subcortical auditory and audiovisual processing of speech and music. Proceedings of the National Academy of Sciences. 104(40):15894–15898.

Nitsche M, Paulus W. 2001. Sustained excitability elevations induced by transcranial DC motor cortex stimulation in humans. Neurology. 57(10):1899–1901.

Oldfield RC. 1971. The assessment and analysis of handedness: the Edinburgh inventory. Neuropsychologia. 9(1):97–113.

Oxenham AJ, Micheyl C, Keebler MV. 2009. Can temporal fine structure represent the fundamental frequency of unresolved harmonics? The Journal of the Acoustical Society of America. 125(4):2189–2199.

Palomar-Garcia MA, Hernandez M, Olcina G, Adrian-Ventura J, Costumero V, Miro-Padilla A, Villar-Rodriguez E, Avila C. 2020. Auditory and frontal anatomic correlates of pitch discrimination in musicians, non-musicians, and children without musical training. Brain Structure and Function. 225(9):2735–2744.

Parbery-Clark A, Marmel F, Bair J, Kraus N. 2011. What subcortical–cortical relationships tell us about processing speech in noise. European Journal of Neuroscience. 33(3):549–557.

Patterson RD, Uppenkamp S, Johnsrude IS, Griffiths TD. 2002. The processing of temporal pitch and melody information in auditory cortex. Neuron. 36(4):767–776.

Penagos H, Melcher JR, Oxenham AJ. 2004. A neural representation of pitch salience in nonprimary human auditory cortex revealed with functional magnetic resonance imaging. Journal of Neuroscience. 24(30):6810–6815.

Presacco A, Simon JZ, Anderson S. 2016. Evidence of degraded representation of speech in noise, in the aging midbrain and cortex. Journal of Neurophysiology. 116(5):2346–2355.

Price CN, Bidelman GM. 2021. Attention reinforces human corticofugal system to aid speech perception in noise. NeuroImage. 235:118014.

Puschmann S, Uppenkamp S, Kollmeier B, Thiel CM. 2010. Dichotic pitch activates pitch processing centre in Heschl’s gyrus. NeuroImage. 49(2):1641–1649.

Ranieri F, Podda MV, Riccardi E, Frisullo G, Dileone M, Profice P, Pilato F, Di Lazzaro V, Grassi C. 2012. Modulation of LTP at rat hippocampal CA3-CA1 synapses by direct current stimulation. Journal of Neurophysiology. 107(7):1868–1880.

Reato D, Rahman A, Bikson M, Parra LC. 2010. Low-intensity electrical stimulation affects network dynamics by modulating population rate and spike timing. Journal of Neuroscience. 30(45):15067–15079.

Ross B, Picton TW, Pantev C. 2002. Temporal integration in the human auditory cortex as represented by the development of the steady-state magnetic field. Hearing Research. 165:68–84.

Ross B, Herdman AT, Pantev C. 2005. Right hemispheric laterality of human 40 Hz auditory steady-state responses. Cerebral Cortex. 15(12):2029–2039.

Ross B, Tremblay KL, Alain C. 2020. Simultaneous EEG and MEG recordings reveal vocal pitch elicited cortical gamma oscillations in young and older adults. NeuroImage. 204:116253.

Russo NM, Skoe E, Trommer B, Nicol T, Zecker S, Bradlow A, Kraus N. 2008. Deficient brainstem encoding of pitch in children with autism spectrum disorders. Clinical Neurophysiology. 119(8):1720–1731.

Salehinejad M, Kuo M, Nitsche M. 2019. The impact of chronotypes and time of the day on tDCS-induced motor cortex plasticity and cortical excitability. Brain Stimulation. 12(2):421.

Schochat E, Rocha-Muniz CN, Filippini R. 2017. Understanding auditory processing disorder through the FFR. In: Kraus N, Anderson S, White-Schwoch T, Fay RR, Popper AN, eds. The Frequency-Following Response: A Window into Human Communication. Springer, Cham. p 225–250..

Sehm B, Schäfer A, Kipping J, Margulies D, Conde V, Taubert M, Villringer A, Ragert P. 2012. Dynamic modulation of intrinsic functional connectivity by transcranial direct current stimulation. Journal of Neurophysiology. 108(12):3253–3263.

Shah B, Nguyen TT, Madhavan S. 2013. Polarity independent effects of cerebellar tDCS on short term ankle visuomotor learning. Brain Stimulation. 6(6):966–968.

Skoe E, Kraus N. 2010. Auditory brainstem response to complex sounds: a tutorial. Ear and Hearing. 31(3):302.

Skoe E, Chandrasekaran B, Spitzer ER, Wong PC, Kraus N. 2014. Human brainstem plasticity: the interaction of stimulus probability and auditory learning. Neurobiology of Learning and Memory. 109:82–93.

Smalt CJ, Krishnan A, Bidelman GM, Ananthakrishnan S, Gandour JT. 2012. Distortion products and their influence on representation of pitch-relevant information in the human brainstem for unresolved harmonic complex tones. Hearing Research. 292:26–34.

Song JH, Skoe E, Banai K, Kraus N. 2011. Perception of speech in noise: neural correlates. Journal of Cognitive Neuroscience. 23(9):2268–2279.

Stagg CJ, Best JG, Stephenson MC, O’Shea J, Wylezinska M, Kincses ZT, Morris PG, Matthews PM, Johansen-Berg H. 2009. Polarity-sensitive modulation of cortical neurotransmitters by transcranial stimulation. Journal of Neuroscience. 29(16):5202–5206.

Strait DL, Kraus N, Skoe E, Ashley R. 2009. Musical experience and neural efficiency: effects of training on subcortical processing of vocal expressions of emotion. European Journal of Neuroscience. 29:661–668.

Suga N. 2020. Plasticity of the adult auditory system based on corticocortical and corticofugal modulations. Neuroscience and Biobehavioral Reviews. 113:461–478.

Suga N, Gao E, Zhang Y, Ma X, Olsen JF. 2000. The corticofugal system for hearing: recent progress. Proceedings of the National Academy of Sciences. 97(22):11807–11814.

Schwenzer M, Mathiak K. 2011. Numeric aspects in pitch identification: an fMRI study. BMC Neuroscience. 12(1):1–9.

Unal G, Bikson M. 2018. Transcranial Direct Current Stimulation (tDCS). In: Krames E, Peckham PH, Rezai AR, eds. Neuromodulation: Comprehensive Textbook of Principles, Technologies, and Therapies. Academic Press. p 1589–1610.

Vanvooren S, Hofmann M, Poelmans H, Ghesquière P, Wouters J. 2015. Theta, beta and gamma rate modulations in the developing auditory system. Hearing Research. 327:153–162.

White-Schwoch T, Carr KW, Thompson EC, Anderson S, Nicol T, Bradlow AR, Zecker SG, Kraus N. 2015. Auditory processing in noise: A preschool biomarker for literacy. PLoS Biology. 13(7):e1002196

Willems RM, Van der Haegen L, Fisher SE, Francks C. 2014. On the other hand: including left-handers in cognitive neuroscience and neurogenetics. Nature Reviews Neuroscience. 15(3):193–201.

Wong PC, Skoe E, Russo NM, Dees T, Kraus N. 2007. Musical experience shapes human brainstem encoding of linguistic pitch patterns. Nature Neuroscience. 10(4):420–422.

Wouters J, McDermott HJ, Francart T. 2015. Sound coding in cochlear implants: From electric pulses to hearing. IEEE Signal Processing Magazine. 32(2):67–80.

Yan, J, Suga N. 1999. Corticofugal amplification of facilitative auditory responses of subcortical combination-sensitive neurons in the mustached bat. Journal of Neurophysiology. 81(2):817–824.

Zaehle T, Beretta M, Jancke L, Herrmann CS, Sandmann P. 2011. Excitability changes induced in the human auditory cortex by transcranial direct current stimulation: Direct electrophysiological evidence. Experimental Brain Research. 215:135–140.

Zatorre RJ, Belin P. 2001. Spectral and temporal processing in human auditory cortex. Cerebral Cortex. 11(10):946–953.

Zatorre RJ, Belin P, Penhune VB. 2002. Structure and function of auditory cortex: music and speech. Trends in Cognitive Sciences. 6(1):37–46.

Zatorre RJ, Evans AC, Meyer E, Gjedde A. 1992. Lateralization of phonetic and pitch discrimination in speech processing. Science. 256(5058):846–849.

Zhang Y, Suga N. 1997. Corticofugal amplification of subcortical responses to single tone stimuli in the mustached bat. Journal of Neurophysiology. 78(6):3489–3492.

